# An RNA foundation model enables discovery of disease mechanisms and candidate therapeutics

**DOI:** 10.1101/2023.09.20.558508

**Authors:** Albi Celaj, Alice Jiexin Gao, Tammy T.Y. Lau, Erle M. Holgersen, Alston Lo, Varun Lodaya, Christopher B. Cole, Robert E. Denroche, Carl Spickett, Omar Wagih, Pedro O. Pinheiro, Parth Vora, Pedrum Mohammadi-Shemirani, Steve Chan, Zach Nussbaum, Xi Zhang, Helen Zhu, Easwaran Ramamurthy, Bhargav Kanuparthi, Michael Iacocca, Diane Ly, Ken Kron, Marta Verby, Kahlin Cheung-Ong, Zvi Shalev, Brandon Vaz, Sakshi Bhargava, Farhan Yusuf, Sharon Samuel, Sabriyeh Alibai, Zahra Baghestani, Xinwen He, Kirsten Krastel, Oladipo Oladapo, Amrudha Mohan, Arathi Shanavas, Magdalena Bugno, Jovanka Bogojeski, Frank Schmitges, Carolyn Kim, Solomon Grant, Rachana Jayaraman, Tehmina Masud, Amit Deshwar, Shreshth Gandhi, Brendan J. Frey

**Affiliations:** Deep Genomics, Toronto, Canada

## Abstract

Accurately modeling and predicting RNA biology has been a long-standing challenge, bearing significant clinical ramifications for variant interpretation and the formulation of tailored therapeutics. We describe a foundation model for RNA biology, “BigRNA”, which was trained on thousands of genome-matched datasets to predict tissue-specific RNA expression, splicing, microRNA sites, and RNA binding protein specificity from DNA sequence. Unlike approaches that are restricted to missense variants, BigRNA can identify pathogenic non-coding variant effects across diverse mechanisms, including polyadenylation, exon skipping and intron retention. BigRNA accurately predicted the effects of steric blocking oligonucleotides (SBOs) on increasing the expression of 4 out of 4 genes, and on splicing for 18 out of 18 exons across 14 genes, including those involved in Wilson disease and spinal muscular atrophy. We anticipate that BigRNA and foundation models like it will have widespread applications in the field of personalized RNA therapeutics.

## Main

Building machine learning models that can predict gene expression from DNA sequence has been a long-standing research goal^1^, and one that has seen significant strides owing to recent advancements in deep learning^2^. These models could revolutionize drug discovery by pinpointing how pathogenic genetic variants alter gene expression and gene processing, and by designing customized drug candidates to counteract these effects^3^. Currently, most efforts have focused on predicting data that measures overall gene expression levels^2,4^, which are not suited to predicting regulatory interventions, for example, specific transcriptional perturbations on splicing or polyadenylation.

RNA sequencing (RNA-seq) data provides a widely-available resource for measuring RNA expression at high resolution and capturing complex transcriptional regulation events across diverse genotypes. This includes both exome variation inherently coded within RNA-seq data itself, and through extensive resources like the Genotype-Tissue Expression^5^ (GTEx) project that pairs RNA-seq with Whole Genome Sequencing (WGS). While building deep neural networks that directly learn from RNA-seq offers the opportunity to understand how changes in DNA sequence lead to changes in complex transcriptional phenotypes, this goal has remained elusive.

We introduce “BigRNA”, a deep learning model that is directly trained on RNA-seq datasets. BigRNA learns from paired genotype and 128bp resolution RNA expression data from many individuals, and can also be applied in a range of downstream tasks such as predicting RNA-binding protein (RBP) specificity and microRNA binding sites. Because BigRNA directly models RNA-seq data, it can discover a diverse set of pathogenic non-coding mechanisms that would each require a specialized model, and can pinpoint their effects on a transcript. We show that BigRNA can discover the effects of non-coding variants on expression and splicing, and matches or exceeds the performance of specialized models in recovering known pathogenic variants.

BigRNA can also help design different types of RNA based therapeutics, including steric blocking oligonucleotides (SBOs). Without any additional training, BigRNA accurately identifies compounds that induce a targeted splicing change, and recovers known approved SBO therapies with high specificity. The ability of BigRNA to understand regulatory mechanisms also allows it to design SBOs that block predicted inhibitory regions to increase the expression of a disease gene. BigRNA represents a new generation of massive deep learning models that can be applied to a range of different personalized RNA therapeutic discovery tasks.

## Results

### BigRNA accurately predicts tissue-specific RNA expression and the binding sites of proteins and microRNAs

To train BigRNA to predict RNA-seq data from the corresponding DNA sequence, we employed a transformer-based architecture^2^ and utilized the GTEx^5^ resource (**Methods**). Given an individual’s genotype, we input two potential haplotypes independently into identical instances of the model, and train it to predict the observed RNA-seq data as the combined output from these haplotypes (**Fig. 1a**, **Supplementary Figs. S1 and S2**). Each output “head” of the model predicts the expression of a single GTEx sample, so that it learns to predict the outputs of 2,956 RNA-seq samples from 70 individuals, covering 51 tissues in total. After training on these RNA-seq datasets, the model is fine-tuned to predict the specificity of RBP and microRNA binding sites (**Fig. 1a**).

**Figure 1.**
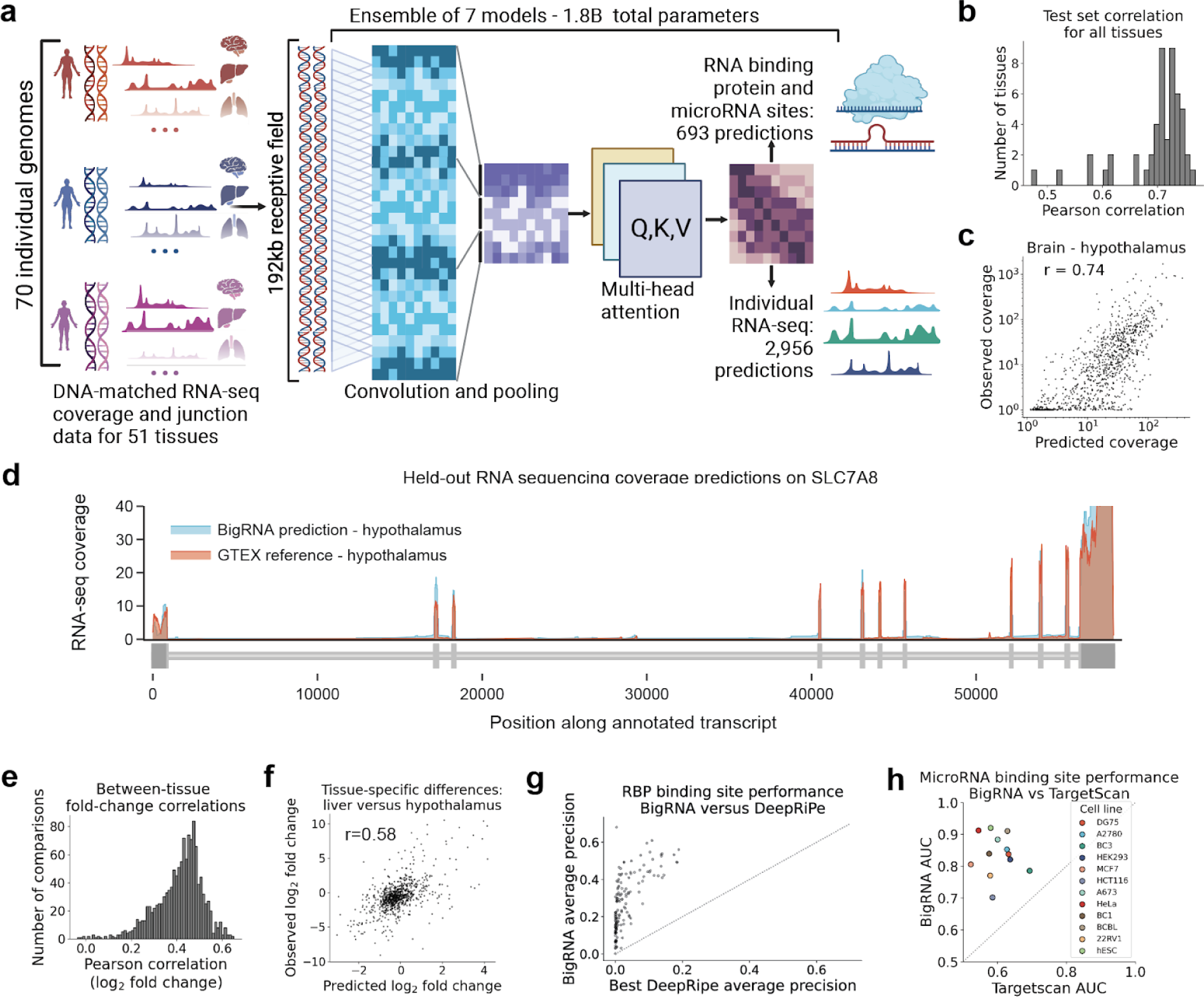
BigRNA accurately predicts tissue-specific RNA expression of unseen sequences. **a.** A schematic of BigRNA’s training. BigRNA was trained on the genomes of 70 individuals, to predict a total of 2,956 RNA-seq datasets over 51 tissues, plus 693 datasets corresponding to RNA binding protein and microRNA sites. **b.** Distribution of correlations between predicted and measured RNA-seq coverage in exonic regions for genes held-out during training (averaged across individuals). **c.** Correlation between predicted and measured RNA-seq coverage for the hypothalamus samples. **d.** Predicted versus measured coverage for *SLC7A8*, averaged across hypothalamus samples for all individuals. **e.** Distribution of correlations between predicted and measured fold-change (pearson r) for all pairwise comparisons across 51 tissues. **f.** Fold-change in gene coverage between liver and hypothalamus. **g.** Comparison of BigRNA and a previously published method, DeepRiPe, for predicting the binding sites of 98 RNA binding proteins across 2 cell lines (142 total experiments). **h.** Comparison of BigRNA and a previously published method, TargetScan, for predicting microRNA binding sites for 12 cell lines.

We first evaluated the ability of BigRNA to predict the expression of unseen genomic sequences. We measured the model’s ability to predict tissue-specific expression levels for all genes outside of genomic regions in the training set. BigRNA exhibited strong performance for predicting expression levels of unseen genes, achieving a correlation coefficient (r) between 0.47 and 0.77 across all tissues (mean=0.70, **Fig. 1b**). We observed slightly stronger performance in brain tissues than non-brain tissues (mean r=0.74 versus 0.69, p=5e-03), and highlight that the model is able to accurately predict expression levels in the hypothalamus (r=0.74, **Fig. 1c**). The ability to predict overall expression levels and capability to accurately delineate intron/exon junctions is illustrated by BigRNA’s predictions for *SLC7A8*, an amino acid transporter within the test set (**Fig. 1d**). To evaluate BigRNA on the much harder task of predicting differences between pairs of tissues, we used BigRNA predictions to compute the fold-change in total exonic coverage between tissue pairs and compared that to observed fold-changes. Across all inter-tissue comparisons, we observed a mean correlation of r=0.4, owing to the increased difficulty of this task (**Fig. 1e**). We highlight a comparison between liver and the hypothalamus (r=0.58, p=7e-64, **Fig. 1f**) to illustrate this capability.

Since drug discovery tasks benefit from clarity of mechanisms, we next examined how well the fine-tuned BigRNA model could predict RBP binding specificity and microRNA binding sites. For the RBP task, we used a large-scale resource of transcriptome-wide binding profiles for 223 datasets covering 150 unique human RBPs in K562 and HepG2 cells^6^. We found that BigRNA achieved high average precision for many RBPs and performed better than the previously-published DeepRiPe^7^ system for all 142 datasets that they had in common (**Fig. 1g**). On predicting microRNA binding sites, BigRNA achieved a median AUC of 0.84 and for all 12 cell lines that we tested, performed better than a previously published method, TargetScan^8^ (**Fig. 1h**). These predictions are useful for identifying regulatory factors that are altered by variants and SBOs (see below).

### Predicting the effects of variants on gene expression

A key challenge in human genetics is to predict the impact of sequence variants that may be found within the human population. Many deep learning models that do well on unseen genes using certain metrics, such as AlphaFold^9^, struggle to predict variant effects^10^. While some accurate methods exist for predicting the pathogenic impact of rare missense variants^11,12^, non-coding variants, such as those located within the 3’ and 5’ untranslated regions (UTRs) of genes, remain difficult to interpret.

To address this gap, we evaluate BigRNA’s ability to predict the impact of a curated set of pathogenic or likely pathogenic (P/LP) UTR variants from ClinVar^13^. We found that BigRNA exhibited strong performance as a general pathogenicity model for variants in both the 3’ UTR and 5’ UTR (AUC=0.95 and 0.8, **Fig. 2a**) by predicting their effects on the expression of their associated disease genes. The weaker performance in the 5’ UTR may be due to a smaller proportion of P/LP variants that modulate RNA expression (18/47 compared to 16/17 for the 3’ UTR amongst variants with known mechanisms, p=0.046), and a substantial proportion of mechanisms that affect translation (29/47). We further investigated a known pathogenic expression-decreasing variant in the 3’ UTR of *NAA10*^14^ (NM_003491.4:c.*43A>G). This variant is known to cause syndromic X-linked microphthalmia, and reduces expression by disrupting the polyadenylation site (PAS) of the *NAA10* transcript. The BigRNA predictions highlight the expression-decreasing effects of this variant (false positive rate, FPR <0.5%), and also predicted the expected lengthening of the 3’ UTR that was observed in RNA-seq samples of affected patients^14^ (**Fig. 2b**). An *in-silico* saturation mutagenesis near this variant highlighted the importance of the PAS, and confirmed the effects of two other nearby P/LP variants (c.*39A>G, c.*40A>G)^14^ (**Fig. 2c**).

**Figure 2.**
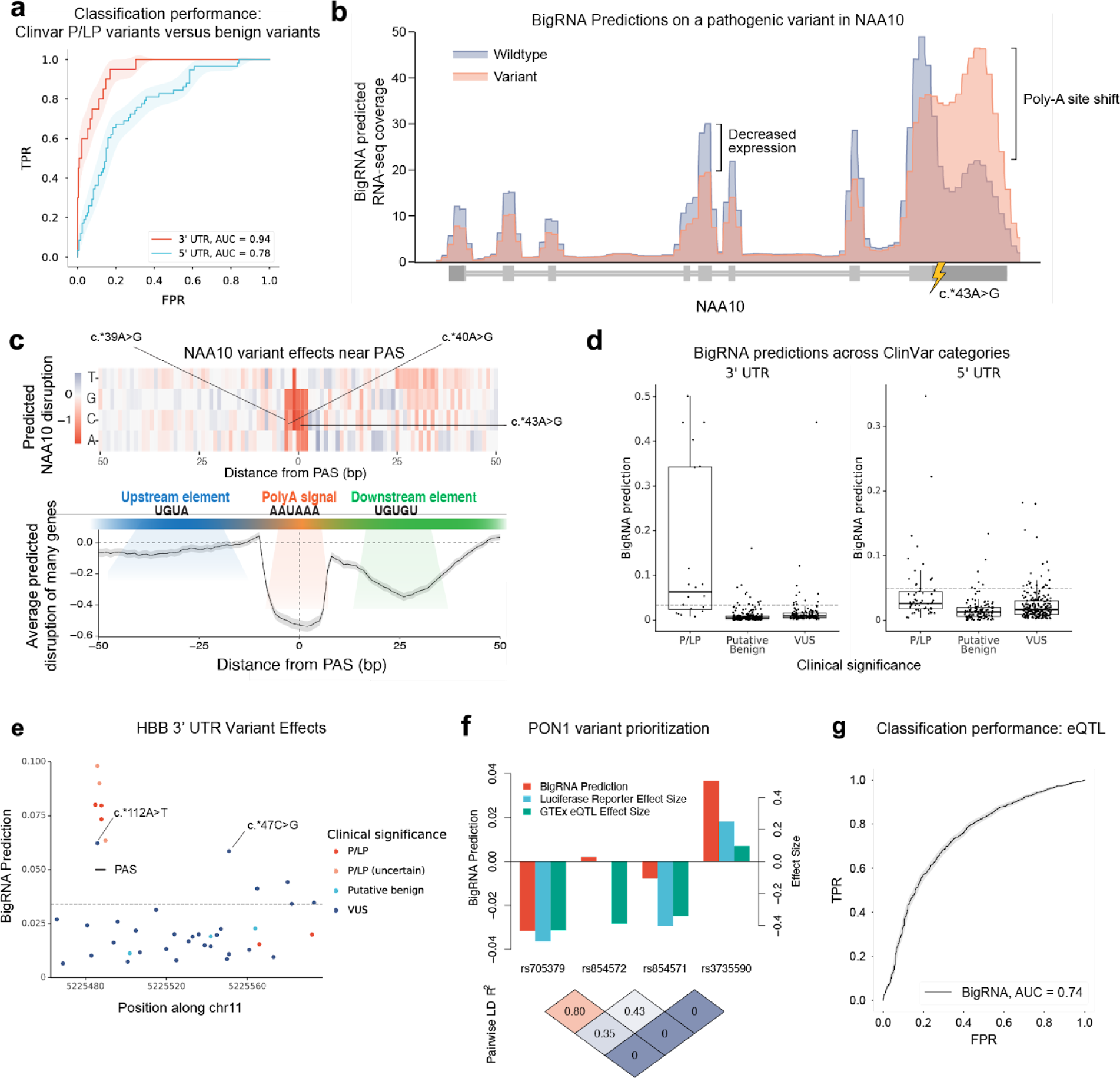
BigRNA predicts the effects of pathogenic expression-modulating variants. **a.** Performance of BigRNA on classifying P/LP variants from putative benign variants in the 3’ UTR and 5’ UTR. **b.** RNA-seq coverage predictions for the effects of a pathogenic variant in the 3’ UTR of NAA10 (NM_003491.4), averaged across all individuals and all tissue types. **c.** Top: BigRNA predictions showing the change in expression for all possible point mutations around the polyadenylation site (PAS) of *NAA10*. Three variants previously identified as impacting the PAS are labeled. Bottom: Relationship between the change in expression predicted by BigRNA from ablating regions around the PAS relative to the distance from the PAS for 200 human poly(A) signal sequences selected from PolyASite 2.0. **d.** The distribution of BigRNA scores for P/LP variants, putative benign variants. and VUS variants from ClinVar for genes included in the UTR benchmarks. The dashed line in both plots (left, y = 0.0341; right, y = 0.0494) represents the threshold of classifying P/LP from putative benign variants at an FPR of 5% in each of the benchmark datasets. **e.** BigRNA predictions for variants of varying clinical significance in *HBB*. The dashed line represents the threshold of classifying P/LP from putative benign variants at a 5% FPR in the 3’ UTR (y = 0.0341). The two highest scoring VUS variants in this gene are annotated. **f.** Top: Comparing BigRNA predicted effects to GTEx eQTL effect size and results of a luciferase reporter assay for four variants suspected to impact PON1 expression. Bottom: Estimated linkage disequilibrium between variants. **g**. Performance of BigRNA at distinguishing fine-mapped expression quantitative trait loci (eQTLs) from controls matched by effector gene (eGene), distance to the transcription start site of the eGene, and minor allele frequency.

We compared BigRNA to Framepool^15^, a ribosomal load model, Saluki^16^, an RNA stability model, and Enformer^2^, an expression model that learns from CAGE-seq. We observed improved performance compared to Enformer for pathogenic variants in both the 5’ and 3’ UTR (p=0.04 and p=0.02, respectively, **Supplementary Fig. S3**). Framepool, a model that predicts ribosomal load^15^, performed similarly to BigRNA for pathogenic variant classification in the 5’ UTR (AUC=0.67 versus 0.78 for BigRNA, p=0.07, **Supplementary Fig. S3**), but BigRNA performed better at classifying the subset of pathogenic 5’ UTR variants that are known to modulate RNA expression (AUC=0.61 versus 0.86 for BigRNA, p=0.002, **Supplementary Fig. S4**). Saluki, an RNA half-life model, had similar performance on the 3’ UTR task (AUC=0.87 vs 0.94 for BigRNA, p=0.27).

Within these genes, we noted many variants of uncertain significance (VUS) in their untranslated regions. Applying BigRNA to these variants at a 5% FPR yielded 12 potential expression-modulating variants in the 3’ UTR (out a total of 139) and 23 in the 5’ UTR (out a total of 222) (**Fig. 2d**). For example, the 3’ UTR of *HBB* had the highest number of VUSs surpassing this threshold (n=6). The highest scoring VUS (NM_000518.5(HBB):c.*112A>T) is in the PAS of this gene, and shares the same position as a known pathogenic variant (c.*112A>G). The PAS region of *HBB* also contains the majority of known P/LP variants (6 of 8). The second-highest scoring VUS (c.*47C>G) was outside of the PAS, and less is known about its function. Looking further, we found that despite being classified as a VUS, this variant is reported to cause decreased expression of *HBB*, supporting the BigRNA prediction^17^. We also noted that three additional P/LP variants in the *HBB* PAS, which were not included in our benchmark due to a lack of evidence in the ClinVar submission^13^, scored above this threshold (**Fig. 2e**), providing computational support for their P/LP classification.

In more genetically complex diseases, it can be challenging to discover causal expression-modulating variant(s) due to linkage disequilibrium (LD). For example, rs705379 and rs854572 are both annotated as expression quantitative trait loci (eQTLs) for Paraoxonase 1 (*PON1*) in GTEx, but a luciferase reporter assay and statistical fine-mapping of the locus show that only rs705379 has an effect on expression^18,19^, which is consistent with BigRNA’s prediction of a much stronger effect and its direction, despite the strong LD. BigRNA also assigned a stronger effect, and correct direction, for two other known expression modulation variants, rs854571 and rs3735590^20^ (**Fig. 2f**). To benchmark BigRNA more broadly, we evaluated its ability to identify fine-mapped eQTLs from negative controls matched on effector gene (eGene), distance to transcription start site (TSS), and minor allele frequency. We saw considerable performance for this task (AUC = 0.74, **Fig. 2g**), improving over Enformer (AUC = 0.70, p=4.8e-04 for difference, **Supplementary Fig. S5**). We note that a series of improvements in eQTL scoring, including matching the predictions to the eQTL tissue of interest, and evaluating over the entire contiguous coding sequence rather than the transcription start site made significant improvements to our performance for both models (**Supplemental Note 1**). BigRNA’s classification performance was similarly strong for variants more than 10 kilobases from their eGene’s TSS (AUC 0.73, versus 0.66 for Enformer, p=8.0e-05 for difference, **Supplementary Fig. S6**). Together, these results indicate that BigRNA is able to help prioritize causal variants that mediate more common diseases, which has been challenging for sequence-based deep neural networks^18,19^.

### Predicting the effects of variants on splicing and intron retention

An important subset of pathogenic variants affect splicing, such as those which cause skipping of an exon. These variants often occur in coding regions, and may be incorrectly classified as benign mutations based on their amino acid substitutions, despite their pathogenic splicing effects^21^. We evaluated BigRNA’s ability to classify the splicing impact of exonic variants that cause substantial (>50%) exon skipping, versus those that do not cause any splicing changes, using results from a massively parallel splicing assay (MaPSy)^21^. By predicting a change in junction coverage caused by these variants, BigRNA was able to accurately predict these skipping variants (AUC = 0.89 **Fig. 3a**), and showed better performance compared to a previously published method, SpliceAI^22^ on this task (AUC=0.80, p<1e-05 for difference, **Supplementary Fig. S7**). We further investigated a pathogenic variant that causes skipping of exon 6 in the *ACADM* gene, leading to a potentially fatal medium-chain acyl-CoA dehydrogenase deficiency^23,24^. BigRNA predicted the exon skipping effects of this variant (FPR = 0.002, **Fig. 3b**), and that it causes this skipping by creating a binding site for the TDP-43 protein^23^, yielding insight into the mechanism-of-action. We further investigated a VUS in *ATP7B* (c.3243+5G>A), a gene which clears copper from liver cells and causes Wilson disease when it is defective^25^. This variant was predicted by BigRNA to cause in-frame skipping of *ATP7B* exon 14 (FPR=0.004, **Fig. 3c**), which contains the ATP site and other critical elements^26^, thus causing a pathogenic loss-of-function. We generated a homozygous HepG2 line and used RT-PCR to assay the effects of this variant and confirm the exon skipping predicted by BigRNA (**Fig. 3c**).

**Figure 3.**
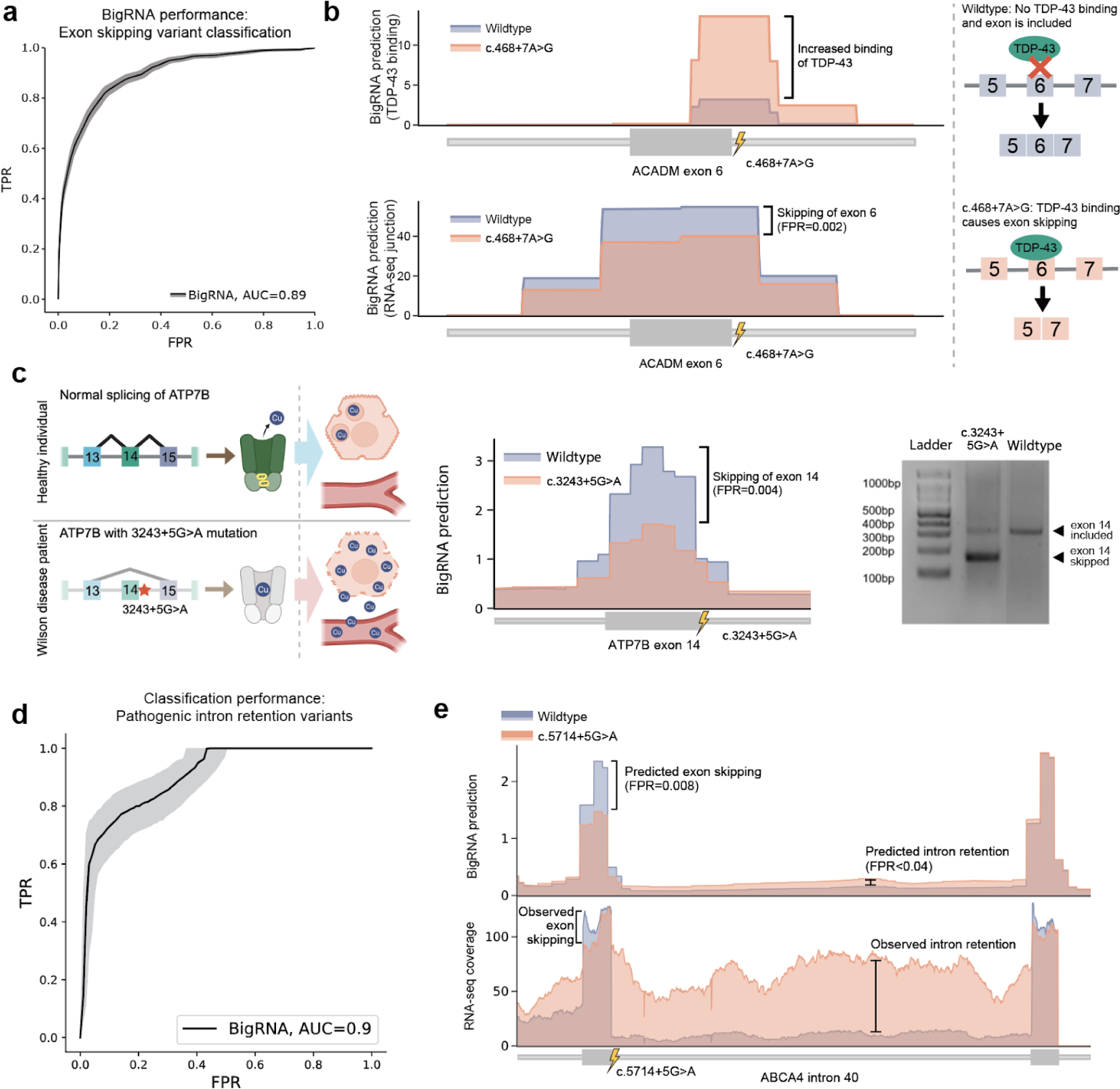
BigRNA captures the effect of variants on splicing. **a.** BigRNA performance on classifying exonic variants that result in exon skipping by at least 50%, from exonic variants that do not cause skipping, both obtained from MaPSy. **b.** BigRNA predicts that the c.468+7A>G variant will result in increased TDP-43 binding and skipping of *ACADM* exon 6. **c.** The *ATP7B* VUS c.3243+5G>A is predicted by BigRNA to cause in-frame skipping of exon 14. This results in reduced levels of functional ATP7B protein, leading to copper buildup in the cell. Right: An RT-PCR in HepG2 cells edited to be homozygous for c.3243+5G>A confirms the expected fragment from exon skipping. **d.** BigRNA performance on classifying variants that cause intron retention (n = 25) from a set of matched variants that do not impact splicing (n = 63). **e.** Top: BigRNA coverage predictions of the c.5714+5G>A variant in *ABCA4*. Bottom: RNA-seq of wildtype WERI cells and WERI cells edited to be homozygous for the variant confirm both exon skipping and intron retention effects.

Another class of pathogenic splicing variants are cryptic splicing mutations that cause full intron retention. We evaluated BigRNA on its ability to predict a set of reported intron retention variants^27^, using nearby common variants as the negative set. We observed strong performance on classifying these mutations (AUC=0.9, **Fig. 3d** and **Supplementary Fig. S8**), so we next investigated whether BigRNA could predict more complex splicing aberrations. We focused our attention on a pathogenic non-canonical splice site variant in the *ABCA4* gene (c.5714+5G>A), which had been found to induce Stargardt disease by causing skipping of *ABCA4* exon 40^28^. This variant was strongly predicted to cause both the skipping of exon 40, and retention of intron 40 (FPR=0.008 and <0.04, respectively, **Fig. 3e**), but the latter had not been reported, likely due to technical limitations in the assay^28^. To test this prediction, we edited a retinoblastoma cell line (WERI-Rb-1) to be homozygous for c.5714+5G>A, and performed RNA sequencing to capture the full suite of splicing events. This confirmed BigRNA’s predictions that this variant causes a complex set of aberrations that includes partial skipping of exon 40, as well as retention of intron 40.

### Designing splice-switching and expression-increase molecules

The ability of BigRNA to understand regulatory mechanisms affecting splicing and gene expression may allow it to design therapeutic interventions that rescue pathogenic variant effects. For this application, we evaluated whether BigRNA could reverse splicing defects by designing steric blocking oligonucleotides (SBOs) – short, chemically-modified synthetic nucleic acid strands purposed to bind specific RNA targets to modulate splicing and gene expression. For example, Nusinersen, an FDA approved SBO, treats spinal muscular atrophy by reversing the skipping of exon 7 in *SMN2*^29^, thus restoring SMN protein levels and mitigating motor neuron loss and muscular atrophy. One way to predict the effect of an SBO is to hide the complementary binding site from the model’s input (**Methods**). This approach is an instance of ‘zero-shot learning’, because no additional task-specific SBO data is used when making the prediction.

To evaluate the utility of zero-shot learning for virtual screening, we first evaluated the ability of BigRNA to re-discover Nusinersen amongst the set of all possible SBOs within 200 base-pairs of *SMN2* exon 7. Strikingly, BigRNA ranked Nusinersen within the top 3 of 437 compounds (**Fig. 4a**). To more systematically evaluate the effectiveness of this approach, we treated 15 exons in 12 genes with a total of 620 SBOs, and observed a strong and statistically significant correlation with the predicted and experimentally-measured exon inclusion levels in all cases (r=0.41-0.77, p=7e-12 to 2e-2, **Fig. 4b**). For comparison, SpliceAI correlated with experiments in 11/15 exons and the correlation was lower than BigRNA for 13/15 exons.

**Figure 4.**
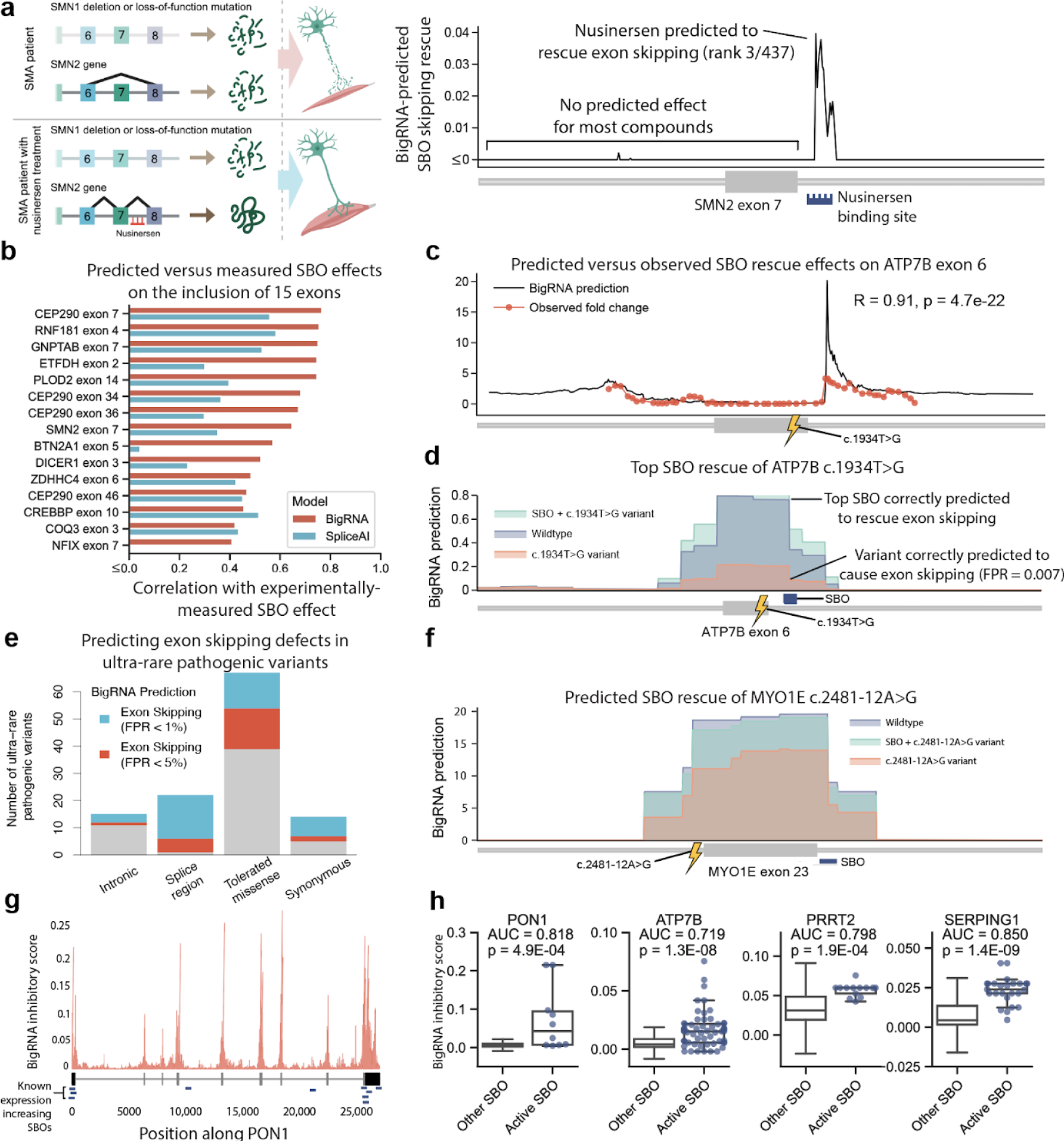
BigRNA predicts the effects of steric blocking oligonucleotides. **a.** Mechanism of action of the splice-switching oligonucleotide Nusinersen, an approved treatment for spinal muscular atrophy (SMA). BigRNA predictions are shown for the exon-restoring effects of all 18-mer SBOs within 200 bp of *SMN2* exon 7. The blue bar shows the position of Nusinersen. Predictions were truncated at zero for the plot. **b.** Spearman correlation between experimentally observed exon-inclusion levels and predictions generated by BigRNA and SpliceAI. A negative correlation for *NFIX* exon 7 versus SpliceAI (r=-0.13) was truncated to zero. **c.** BigRNA predictions of SBO effects on *ATP7B* exon 6 inclusion. 55 SBOs were screened by qPCR to measure total *ATP7B* expression relative to control (fold change), and the Spearman correlation was computed between the BigRNA predictions and observed fold changes. **d.** BigRNA predictions for wildtype, Met645Arg (c.1934T>G) variant, and Met645Arg variant with treatment (lead SBO targeting *ATP7B* exon 6). The junction count tracks pertaining to individual samples of the liver tissue are averaged for plotting. **e.** Proportion of ultra-rare pathogenic variants associated with AR disorders with BigRNA exon skipping predictions above the 1% and 5% FPR thresholds. Intronic (>8bp from splice site), splice region (<8bp from splice site excluding the core dinucleotides), tolerated missense (SIFT score > 0.05) and synonymous variants are shown. **f.** BigRNA predictions for wildtype, c.2481-12A>G variant and the variant with treatment (lead SBO targeting *MYO1E* exon 23). **g.** BigRNA predicts expression increase SBOs in *PON1*. BigRNA inhibitory scores are plotted by region of the gene. The transcript structure is shown under the scores, and the locations of the 10 dose-response hits are shown with blue bars. The distribution of BigRNA inhibitory scores for the 10 dose-response hits is significantly different from the distribution for other length-matched SBOs targeting *PON1* **h.** BigRNA scores of screening hits compared to background of all possible SBOs of same length for *PON1, ATP7B*, *PRRT2*, and *SERPING1*.

We then used BigRNA to design a novel splice-switching SBO that rescues a pathogenic splicing defect. Previously, we had reported that a missense variant in the *ATP7B* gene (c.1934T>G, Met645Arg) leads to Wilson disease by promoting skipping of exon 6, thus resulting in lowered levels of functional protein and subsequent copper accumulation in liver cells^25^. We created a disease model of the Met645Arg variant in HepG2 cells, and used this system to test a set of SBOs targeting the skipped exon (**Methods**). We observed a strong relationship between the predicted and measured splicing changes (r=0.91, p=4.7e-22, **Fig. 4c**). The top compound from this assay was predicted to be in the top 7 of 458 possible compounds by BigRNA. To summarize, BigRNA predicted both the exon skipping caused by Met645Arg (FPR=0.007) and the restorative effect of the top experimentally-validated compound (**Fig. 4d**).

BigRNA’s ability to score SBOs has utility in developing therapeutic candidates targeting extremely rare variants within a constrained budget. First, we evaluated BigRNA’s ability to score SBOs that target a pseudo-exon in the *ATM* gene caused by the rare c.5763-1050A>G mutation, leading to ataxia-telangiectasia^30^. We observed significant correlation between the predictions and experimentally observed splicing efficiencies (r=0.64, p=3.3e-04, **Supplementary Fig. S9**), and ranked the lead therapeutic candidate in the top 7 of 516 possible compounds. We sought to explore whether similar therapeutic candidates could be developed for other rare splicing diseases. After curating a set of extremely rare, so-called “N=1”, pathogenic variants from ClinVar (**Methods**), we used BigRNA to predict which ones are likely to act through exon skipping while not affecting the core splice donor or acceptor site (**Fig. 4e**), thus potentially being eligible for SBO remediation. This included synonymous variants, non-synonymous variants predicted to be tolerated^31^, and variants near splice sites. One such variant was in intron 22 of *MYO1E,* which is associated with glomerulosclerosis^32^. While no published mechanism exists for this variant, it was predicted to cause skipping of exon 23, and the top SBO was predicted to completely rescue this skipping defect, suggesting that this variant is amenable to personalized SBO treatment (**Fig. 4f**).

Owing to BigRNA’s striking ability to help design splice-switching SBOs, we turned to the more challenging problem of designing SBOs that amplify gene expression. This requires the model to rank all possible compounds targeting any part of the transcript, again without any additional training and additionally with no prior knowledge of inhibitory regions. Due to the greatly increased search space, we first developed a method to score a large number of compounds in a computationally efficient manner. For this, we applied a combination of established saliency mapping techniques^33,34^ to evaluate the contribution of each base pair in a transcript on its expression in a given tissue, and took the minimum contribution score at the SBO binding region as the ‘inhibitory score’ of each compound (**Methods**). We again benchmarked this scoring on Nusinersen, reasoning that the skipping of exon 7 and subsequent nonsense-mediated mRNA decay is a major expression bottleneck. Considering all 26,901 SBOs of length 18, Nusinersen ranked in the top 2.28% (**Supplementary Fig. S10**), suggesting that BigRNA’s inhibitory scores can be used to identify inhibitory regions, and that this strategy could have recovered Nusinersen within a tractable screening budget.

We then sought to systematically assess how well BigRNA could be used to discover novel therapeutically beneficial expression-increasing SBOs. An example is Paraoxonase 1 (*PON1*), where variants that decrease expression of the gene or catalytic activity of the protein have been associated with an increased risk of atherosclerotic cardiovascular disease^35,36^ (ASCVD). In murine models, modulation of *PON1* expression has been shown to directionally influence the risk of ASCVD and related phenotypes^37–40^, thus presenting a compelling opportunity for expression-increasing therapeutics. We used BigRNA to perform large-scale SBO design, experimentally tested the predicted SBOs in primary human hepatocytes, and identified 10 compounds that showed activity for increasing *PON1* expression (**Methods**). By using a liver-specific score to rank all positive compounds, BigRNA showed a strong ability to prioritize expression increase compounds (AUC=0.818, **Fig. 4h**). To expand this study, we screened expression-increasing compounds for *ATP7B* (to benefit a broader population beyond Met645Arg), as well as *PRRT2* and *SERPING1*, which may confer therapeutic benefits for benign familial infantile epilepsy^41^ and hereditary angioedema^42^. For all three genes, BigRNA’s predictions successfully prioritized expression-increasing SBOs without requiring any additional training (AUC=0.72-0.85, **Fig. 4h**).

## Discussion

The rapid evolution of computational models in genomics has enabled the use of methods that can learn from large-scale genomics data to predict RNA expression from DNA sequence. Using deep learning to model RNA-seq data and take into account individual genomic sequence variation, we can enable novel and accelerated discovery on several drug discovery tasks.

When we adapted previously published deep learning systems to the drug discovery tasks that we evaluated, we found that BigRNA performed substantially better overall. It improved significantly over specialized models like TargetScan^8^ and DeepRiPe^7^ for predicting microRNA and RBP binding sites, and was more accurate than SpliceAI^22^ at identifying exon skipping variants as well as designing splice-switching SBOs. BigRNA could accurately predict pathogenic variants in untranslated regions, matching specialized models for the 5’ and 3’ UTRs^15,16^, and improved upon the general-purpose Enformer model^2^. In cases where BigRNA’s performance matched an existing model, direct modeling of RNA-seq data had distinct advantages. For example, unlike a previously described ribosomal loading model^15^, BigRNA could predict all classes of pathogenic mutations in the 5’ UTR, and unlike a model of RNA half life^16^, it could predict that a pathogenic variant acts by changing the polyadenylation site, which reduces the half-life. Existing methods for predicting splice donor and acceptor strength^22^ are unable to identify correlated splicing events, such as intron retention, but we found that BigRNA is able to do so. For complex traits, in contrast to traditional fine-mapping methods that do not provide insight into the mechanistic impact of causal mutations^43^, BigRNA can make predictions for complex trait heritability contributions from many different mechanisms that do not exert their effect through a change in protein structure.

The ability of BigRNA to learn mechanisms of RNA regulation is reflected by the fact that it was able to accurately design SBOs that counteract the effects of pathogenic variants or that increase gene expression, without being provided with a single training case of an SBO and its effect. Nonetheless, a further avenue of work would include fine-tuning BigRNA by learning from SBO treatment data, such as from the rich information encoded by SBO-treated RNA-seq samples^44^. Similar approaches can be used for other therapeutic modalities such as predicting the phenotypic effects of induced ADAR (adenosine deaminase acting on RNA) editing so that they confer a similar compensatory effect on splicing or expression^45^, or designing mRNAs that have increased half-life and translation efficiency.

Several avenues exist to improve the predictive abilities of BigRNA. The 128bp resolution of the model can be improved with additional training resources^2^. Improvements in the speed and scalability of the transformer architecture^46^, coupled with the use of parameterized upsampling^47^ may allow the model to retain a high context size while producing predictions at single base-pair resolution. Training on more individuals could improve generalization across genotypes. While the training procedure takes into account variation from 70 individuals, WGS-paired RNA-seq data is available for many more GTEx samples, and can be supplemented with additional datasets^48^. To take into account such a large amount of data, methods have been developed to prioritize the most informative training points^49^, allowing the training procedure to scale and effectively learn from extremely large datasets. To explore improved prediction of differences between individuals, a contrastive training objective can be used^50,51,52^ and predictions can be made for the difference in expression between two haplotypes^53^.

Our results show that different drug discovery tasks can be assisted by deep learning. We believe that BigRNA and deep learning systems like it have the potential to transform the field of RNA therapeutics.

## Methods

### RNA-seq model training

We downloaded and aligned RNA-seq data from the GTEx consortium^5^ V6 release, processing all available data from the set of 70 individuals with the most tissue availability (data from a total of 51 tissues are available, but data availability varies between individuals). Data was processed using an in-house pipeline (Supplementary Information 1.2). Each RNA-seq sample was processed into two data tracks: coverage and junction, where the junction track contains a subset of read counts at splice junctions. To make the data compatible with the 128bp resolution of the model’s architecture^2^, we applied 128bp-window average-pooling on coverage tracks, and 128bp-window sum-pooling on junction tracks. To incorporate genomic variants from each individual, we re-aligned the RNA-seq data to match the insertions and deletions introduced by each individual’s haplotype (Supplementary Information 1.2). BigRNA was trained with a separate output for each sample, so that each output can be independently learned. We trained BigRNA by minimizing differences between prediction from both haplotypes and the observed coverage and junction tracks from RNA-seq (Supplementary Information 1.3, Supplementary Equation S2). In addition to the individual-specific outputs, we also added individual-agnostic per-tissue outputs to encourage the model to learn a mapping from genotype to expected expression (where the expectation is taken across all individuals). Description of all output heads can be found in Supplementary Data 1. Fig. 1a shows the training pipeline. The same procedure was used to train an ensemble of 7 models, varying learning rate, degree of gradient clipping, and the pre-training strategy for each model in the ensemble (Supplementary Information 1.3, Supplementary Table S1). At inference time, to predict on a genomic interval, we used shifted intervals to increase the prediction resolution to 64 base pairs, and averaged predictions from both strands (Supplementary Information 1.4, Supplementary Fig. S1, Supplementary Fig. S2).

### Fine-tuning on RBP and microRNA datƒasets

After training models on RNA-seq dataset, we further fine-tuned models on RBP and microRNA datasets. The RBP dataset was constructed by downloading eCLIP data^6^ from ENCODE^54^ (Supplementary Information 1.2.2). The microRNA dataset was generated by processing CLIP-Seq data from 12 cell lines (Supplementary Information 1.2.3). We fine-tuned the model by first updating weights of the last layer for 10 epochs, then updating weights of the entire model for another 30 epochs (Supplementary Information 1.3). Description of all output heads can be found in Supplementary Data 2.

### Held-out performance on gene expression and differential gene expression

We selected protein coding genes that are completely outside the training and validation set, and which overlap at least one interval in the test set. Predictions and targets were mean-aggregated over all exons for each gene to yield one value per gene (Supplementary Information 2.1). For each tissue, we compute the correlation between prediction and target across all genes. To evaluate performance on differential gene expression, we constructed all pairwise comparisons between tissues, and computed the log_2_ fold-change using the predicted and target coverage data (Supplementary Information 2.2). For each tissue pair we computed the correlation between the predicted and target log_2_ fold-changes across all genes.

### Visualizing prediction on SLC7A8

Sequence of SLC7A8 gene was obtained from hg38 genome build with Gencode v29 annotation. We averaged output heads that correspond to coverage in the “Brain - Hypothalamus” tissue to obtain BigRNA prediction for visualization (Supplementary Information 2.3).

### Held-out performance on RBP

Processed RBP peaks were obtained from ENCODE^54^, and processed into low resolution binary labels by taking into account noise in the data [Supplementary Information 1.2.2, Supplementary Equation S8]. We selected protein coding genes that are completely outside the training and validation set, and made predictions using BigRNA and DeepRiPe^7^. Both BigRNA and DeepRiPe predictions were averaged within each 128-bp window (Supplementary Information 2.4). Fig. 1g shows the average precision performance of BigRNA and DeepRiPe.

### Held-out performance on microRNA

The microRNA dataset was generated by processing CLIP-Seq data from 12 cell lines (Supplementary Information 1.2.3). The called peaks were further processed into low resolution binary labels by taking into account noise in the data (Supplementary Equation S9). We selected protein coding genes that are completely outside the training and validation set, and made predictions using BigRNA and TargetScan^8^. Both BigRNA and TargetScan predictions were averaged within each 128-bp window (Supplementary Information 2.5). Fig 1h shows the au-ROC performance of BigRNA and TargetScan.

### Benchmarking variant effect predictions on pathogenic variants

Pathogenic or likely pathogenic (P/LP) UTR SNVs were obtained from Bohn et al^13^. Putative benign SNVs located in the same UTR were obtained from ClinVar, if they were classified as benign or likely benign (B/LB), and gnomAD v3 if their global allele frequency was greater than 0.001^55^ (Supplementary Information 3.1.1). For the 5’ UTR benchmark, we predicted the effect of the variant using BigRNA, Enformer, and FramePool and took the absolute value of the variant effect scores. For the 3’ UTR benchmark, we evaluated BigRNA, Enformer, and Saluki and again, took the absolute values of the variant effect scores (Supplementary Information 3.1.2, Supplementary Equation S10-12). In addition to Fig. 2d, Supplementary Fig. S3 shows the ROC curve and PRC of classification performance of all models. To compare models, we performed permutation tests with 10000 permutations (Supplementary Information 3.1.3). Variants of uncertain significance (VUS) in the UTRs of the genes that were in the benchmark were extracted as described in Supplementary Information 3.1.4.

### Predicting the impact of disrupting polyadenylation sites

To evaluate BigRNA’s ability to predict poly(A) sites, we conducted an in-silico 11 bp N-mask tiling analysis across each poly(A) region. Poly(A) sites (PAS) from 200 genes were obtained from PolyASite 2.0^56^ (Supplementary Information 3.3). For each PAS, we expanded the site by ±100 bp to cover proximal regulatory elements, resulting in 206 bp regions. We subsequently N-masked 11 bp tiles across the region and compared BigRNA predictions for the N-masked sequences (mutant) and the poly(A) signal sequence (wildtype). The BigRNA predictions were based on the mean of the individual sample RNA-seq coverage heads across all tissue types (Supplementary Information 3.3). For the *NAA10* PAS and its surrounding 100 bp context, we performed saturation mutagenesis by point-mutating every reference nucleic acid base to every other nucleic acid base. Similar to the poly(A) site analysis, we carried out predictions using the BigRNA model to assess the impact of these mutations on gene expression.

### Expression quantitative trait loci (eQTLs) and linkage disequilibrium (LD) estimation for *PON1* variants

The four variants with known expression effect were rs705379 (chr7:95324583:G:A), rs854571 (chr7:95325307:T:C), rs854572 (chr7:95325384:C:G) and rs3735590 (chr7:95298183:G:A). The eQTL and normalized effect size of these variants on *PON1* liver tissue expression were obtained from the GTEx eQTL Calculator. The LD R^2^ values between variants was calculated using the NIH LDmatrix tool with the GBR population selected.

### Classifying expression quantitative trait loci (eQTLs) versus matched controls

To construct a benchmark dataset from confidently fine-mapped eQTLs, variants with a posterior inclusion probability of 0.5 or greater (indicating that they are the most likely causal variant in the credible set) were selected from eQTLGen statistical fine-mapping of expression modulating variants in GTEx v8^19^. eQTLs within 50kbp of the transcription start site of the primary or most highly expressed transcript for the reported eGene were selected to ensure that deep learning models would have sufficient genomic context to accurately predict changes in expression. For each eQTL we selected a matched negative control variant from the same effector gene (eGene) which was not associated with its expression (P > 0.05) in any tissue and within 10% of the eQTL’s minor allele frequency and 10kbp of the eQTL’s genomic position. This resulted in a dataset of 1374 eQTL variants and 1162 matched negative controls.

### Classifying variants that cause intron retention

Variants that cause full intron retention were manually curated from splicing variants downloaded from the SPCards database^27^. A matching set of variants that do not cause intron retention were processed from gnomAD^55^ (Supplementary Information 4.1.1). For each variant, we use BigRNA to predict the relative coverage between intron and the two flanking exons, and compute the score as the ratio between wild-type and mutant-type, aggregated across models in ensemble (Supplementary Information 4.1.2, Supplementary Equation S14-16).

### Classifying variants that cause exon skipping

For each mutation in the MaPSy dataset^21^, we computed the splicing odds ratio and confidence interval using the reported readout from both in-vitro and in-vivo assays, to create a high confidence binary label on skipping versus non-skipping at splicing levels ranging from 50% to 10% (Supplementary Information 4.2.1, Supplementary Equation S17-18). For each mutation, we used BigRNA to predict the difference in junction counts between wild-type and mutant-type, normalized by exon, and aggregate across models in ensemble (Supplementary Information 4.2.2, Supplementary Equation S19). Fig 3a shows ROC curve of classification performance on skipping versus non-skipping at 50% splicing level. For model comparison (Supplementary Fig. S7), we performed permutation tests with 100,000 permutations.

### Predicting the effect of splice-switching SBOs

To obtain the relative ranking of Nusinersen, we ranked all possible SBOs of length 18 within 200 base pairs of exon 7 of SMN2 (Supplementary Information 5.1). We used RT-PCR to measure the Percentage Spliced In (PSI) values for 15 exons in the HEK293T cell line, and compared the measured PSI with the predicted SBO effect of SpliceAI and BigRNA using the Spearman Correlation metric (Supplementary Information 5.2). We repeated the above evaluation for SBOs targeting Met645Arg; here we edited HepG2 cells to introduce the c.1934T>G Met645Arg variant, and screened a library of 55 SBOs by qPCR. Spearman correlation was computed between BigRNA predictions and the experimentally observed *ATP7B* expression levels (Supplementary Information 5.3). The same evaluation was carried out on published data of SBOs designed to skip a pseudo-exon created by the c.5763-1050A>G variant in ATM^30^.

The set of “N=1” variants was created by selecting pathogenic or likely pathogenic variants (ClinVar) from genes that are exclusively associated with autosomal recessive disorders (OMIM). BigRNA predictions were made for SNVs with very low estimated worldwide prevalence (n=1582, GnomAD) and we curated synonymous, tolerated missense (SIFT) and intronic variants (excluding the core dinucleotides) for their mechanisms of pathogenicity (Supplementary Information 5.5). All possible 20-mers within 200 bp of *MYO1E* exon 23 were scored for their ability to remedy the effect of the c.2481-12A>G variant, and we visualized the predictions for the highest ranked SBO.

### Predicting the effect of expression increase SBOs

Expression increase can occur through a variety of mechanisms, and SBOs can be designed anywhere in the gene. By applying a combination of established saliency mapping techniques^33,34^, we evaluated the contribution of each base pair in a transcript to the expression of the related gene in the relevant tissue, yielding a sensitivity score for each base pair’s impact on gene expression levels, called the Inhibitory Score (Supplementary Information 1.5). This per-base-pair score was then used to rank SBOs by taking the minimum score of any overlapping base-pair (Supplementary Information 5.6.2). For the Nusinersen ranking evaluation, we used the BigRNA Inhibitory Score to score all candidate SBOs of length 18 targeting the entirety of the gene body of SMN2. The same process was applied to expression increase SBOs identified from screens of *PON1*, *ATPB*, *PRRT2*, and *SERPING1*. Scores between hit SBOs and the background of all candidate SBOs were compared with a Mann-Whitney U-Test.

### Data Availability

Data and code to be made available upon peer review.

## Acknowledgements

We thank David Kelley for advice, Janine Truong for editing figures and reviewing the manuscript, and Laurence MacPhie for reviewing the manuscript. We also thank Daniele Merico for help in conceiving of the appropriate benchmarks, and Tim Yu for suggesting we include results for ultra-rare variants.

## Contributions

A.C., A.J.G., and B.F. initiated the project. A.C., A.J.G., T.T.Y.L., E.M.H., and C.B.C. conceived of the study and designed analyses. A.C., P.O.P. and Z.N. designed the model. A.J.G., A.L., V.L., and S.C. helped implement and improve the model. A.C., X.Z., P.V., and H.Z. processed the training data. A.J.G., T.T.Y.L., E.M.H., A.L., V.L., C.B.C., R.E.D., O.W., P.V., E.R., and B.K. performed benchmarking and downstream analyses. P.V., P.M.S, M.I., E.R., C.K., and S.Gr. aided with benchmarking. E.M.H., V.L., P.V., and R.J. processed experimental data. C.S., D.L., K.K., M.V., K.C.O, Z.S., B.V., S.B., F.Y., S.S., S.A., Z.B., X.H., K.K., O.O., A.M., A.S., M.B., J.B., and F.S. performed validation experiments; T.M., A.D., S.Gh., and B.F. supervised the study

## Ethics Declaration

### Competing interests

All listed authors are present or past employees of Deep Genomics Inc. This study received funding from Deep Genomics in the form of salary support and covering of computational costs. The founder was involved in the decision to submit for publication.

## Supplementary Information

In a separate document.

