## Supplementary Information for "An RNA foundation model enables discovery of disease mechanisms and candidate therapeutics"

**4 Variant effect on splicing** 19
- 4.1 Intron retention 19
  - 4.1.1 Data processing 19
  - 4.1.2 Predictive scores 19
  - 4.1.3 Evaluation 20
- 4.2 Exon skipping - MaPSy evaluation 20
  - 4.2.1 Data processing 20
  - 4.2.2 Predictive scores 21
  - 4.2.3 Evaluation 22
- 4.3 *ACADM* variant effect on TDP-43 binding 22
- 4.4 Validating variant effect of *ATP7B* mutation 22
- 4.5 Validating variant effect of *ABCA4* mutation 23

**5 Designing steric blocking oligonucleotides (SBO)** 24
- 5.1 Scoring SBOs for splicing effects 24
- 5.2 LabChip exon skipping 24
- 5.3 Screening *ATP7B* exon 6 M645R SBOs 24
- 5.4 Visualization of BigRNA predictions for wildtype vs. M645R variant vs. oligonucleotide and M645R variant in *ATP7B* 25
- 5.5 Identification of "N=1" variants associated with autosomal recessive disorders 26
- 5.6 Scoring expression increase SBOs 26
  - 5.6.1 SBO screen for *PON1* 27
  - 5.6.2 SBO screens for *ATP7B*, *PRRT2*, and *SERPING1* 28

**Supplementary Figures** 29

**Supplementary Tables** 38

**Supplementary Notes** 39
- Scoring considerations for eQTL evaluation 39

### 1 Modelling

This section describes the methodological details relevant to **Fig. 1a**.

Our models and training pipelines were implemented in TensorFlow [1].

#### 1.1 Model architecture

We adapted the architecture of Enformer [2] and changed the width of the final head layer to suit our training tasks. The input to BigRNA is a 196608bp DNA sequence that is encoded as a $196608 \times 4$ one-hot matrix using the base labelling scheme $A = 0$, $C = 1$, $G = 2$, and $T = 3$. Its output is a $896 \times d$ matrix of predictions for $d$ output heads, corresponding to the centered $896 \times 128 = 114688$bp subsequence of the input, given at 128bp resolution.

#### 1.2 Datasets

##### 1.2.1 RNA-seq

Each example in our training dataset contains **(1)** a pair of aligned genomic intervals for the two haplotypes of a single individual (out of 70) and **(2)** aligned RNA-seq data for each haplotype, for each tissue type available for that individual. Our full data processing pipeline works as follows:

**RNA-seq processing.** First, we constructed genomic data tracks from the Genotype-Tissue Expression (GTEx) project [3], which contains tissue-specific RNA-seq data from a large number of individuals. This was done in the following steps:

1. **GTEx download.** We downloaded Sequence Read Archive (SRA) files from GTEx. Then, we converted SRA to FASTQ format using SRA Toolkit 2.7.0 [4] by calling: `fastq-dump --split-3`.
2. **Comprehensive splice junction annotation.** To minimize annotation bias[1] in the alignment, we created a comprehensive set of annotation by combining splice junctions from Gencode v25 [5] and Intropolis v1 [6]. To avoid false positives in Intropolis (due to high sequencing depth), we further processed the dataset to those that satisfy the following criteria:
   - Supported by 2 or more samples
   - Supported by 5 or more reads in aggregate
   - At least one end is annotated in Gencode v25 [5]
   - Spliced at least 0.01% of the time (among the junctions that share the same annotated donor/acceptor)
3. **HISAT2 index.** We called `hisat2-build` from HISAT2 2.0.4 [7] to generate the genome annotation index, using human genome build GRCh37 release 75, Gencode v25 [5], and the above mentioned comprehensive splice junction annotation. We also incorporated `dbSNP 146` to build a SNP-aware index.

[1] Annotation bias: preferential alignment to annotated vs. unannotated junction.

4. **Pre-alignment QC.** We used Cutadapt 1.10 [8] to trim adapter and low-quality tail, and used Trimmomatic 0.36 [9] to remove reads with low quality window (average quality score of 2 nt window below 10) as well as reads shorter than 30 nt.
5. **Alignment.** We used HISAT2 2.0.4 [7] to perform a two-pass alignment procedure. Pass 1 aligned the last 40M of the paired end reads of the sample on the genome-annotation index to discover sample-specific novel splice junctions.[2] Pass 2 aligned all reads on the genome-annotation index that is augmented with the novel junctions found in Pass 1.
6. **Post-alignment QC.** We discarded unmapped reads, multi-mapped reads, and reads with edit distance larger than 2.[3] Alignments for paired and unpaired reads were combined. Alignment was then sorted, compressed and indexed as BAM files.
7. **Coverage and junction data.** In addition to the BAM files produced from the above step (which we'll refer to as *coverage* data), for each sample in the dataset, we also extracted all the spliced read alignments, which we'll refer to as *junction* data. To create training data track for coverage data, we used `bedtools genomecov` to generate per-position coverage. To create training data track for junction data, we processed the BAM files using an in-house pipeline, and counted the number of spliced reads for each unique splice junction (either acceptor or donor site).

**Interval selection.** We used the same intervals and training/validation/test split as Enformer [2].

**Haplotype construction.** For each genomic region, we generated multiple training examples, one for each individual. We used genotype data from GTEx [3] to query variants overlapping the region for the particular individual, and constructed the diploid genomic sequence by mutating the reference genomic sequence with the variants. Since we did not have access to phasing information, we partitioned the set of variants randomly[4] into two disjoint sets, and applied each set of variants to mutate the reference genome to get two haplotype sequences, which we called haplotype 0 and haplotype 1. These two sequence can be of different lengths, and required further re-alignment (discussed below).

**RNA-seq data re-alignment.** Since RNA-seq reads were aligned to reference genome, we need to re-align the data to the two haplotypes. Given the above two haplotype sequences, we constructed two position alignments, one between reference genomic sequence positions and haplotype 0 sequence positions, and the other between reference genomic sequence positions and haplotype 1 sequence positions. Given the alignment, we constructed a mapping from positions in reference genomic sequence to the aligned

---

[2] 40M paired-end reads is considered a plenty representative subset of each sample to detect sample-specific splice junctions.

[3] With each soft clipping treated as $\frac{1}{3}$ mismatches.

[4] We partition the set of variants randomly onto each of the haplotype based on the following assumption: if we decompose the total expression of a given transcript as the sum of two "wild-type" sequences, plus individual additive effects of each variant, then the process of adding the output from both variants is invariant to what haplotype each variant is assigned to. However, this procedure does not account for from within-haplotype variant-variant genetic interactions, which requires phased data.

position in each haplotype, with insertion positions repeating data directly upstream of the insertion interval. We applied the above mapping to translate RNA-seq data to align with each of the two haplotypes.

**Intra-haplotype alignment.** After applying the variants, we have two DNA sequences $\mathbf{x}_0$ and $\mathbf{x}_1$, one for each haplotype of individual $1 \leq \ell \leq 70$. However, $\mathbf{x}_0$ and $\mathbf{x}_1$ may be of different lengths or longer than the 196608bp expected by the model, so we need to crop both sequences. For example, a simple strategy is to truncate $\mathbf{x}_i$ to its first 196608 tokens. Once uniformly-sized intervals are chosen, for each haplotype, we can then read off the targets as follows: we collect the $d_\ell$ RNA-seq samples obtained from the desired haplotype of individual $\ell$, which ranges over a subset of 51 possible tissue types (so $d_\ell \leq 51$). Recall that each sample produces two tracks, coverage and junction, so there are a total of $2d_\ell$ tracks for individual $\ell$. We re-scale each track by $C_{50}/C$, where $C$ is the coverage of the track's sample and $C_{50}$ is the median coverage across all samples in GTEx (including those from other individuals). Then, we read off the track data at the center 114688bp of the haplotype sequence and pool it to a 128bp resolution, using a mean-aggregation for coverage data and a sum-aggregation for junction data.[5] After concatenating all samples and repeating the process for both haplotypes, we obtain two $896 \times 2d_\ell$ target matrices $\mathbf{T}_0$ and $\mathbf{T}_1$ for haplotypes $\mathbf{x}_0$ and $\mathbf{x}_1$, respectively.

However, our initial choice of left-cropping both sequences $\mathbf{x}_i$ may be suboptimal, since the haplotypes may be mis-aligned. Because we can only learn the sum of both haplotypes rather than the individual contributions of each (since we do not have allele-specific data), we attempted to align the data together as much as possible given the limitations of the 128bp resolution. For this, we greedily searched for a more optimal cropping by considering progressively shifted crops of one of the haplotype sequences. Formally, for $k$ ranging from 0 to 31, we iteratively considered cropping the first 196608 tokens of one haplotype sequence and the $(k+1)$-th to $(k+196608)$-th tokens of the other. We picked the cropping that maximizes the Pearson correlation $r$ between the entries of the derived haplotype targets $\mathbf{T}_0$ and $\mathbf{T}_1$, but if a sufficiently-high correlation ($r \geq 0.999$) is found at any step, then we terminated our search early.

**Post-processing.** In the preceding step, note that the second dimension of the target matrices varies based on the individual that the haplotype pertains to. A final step is then to pad the target matrices to a uniform shape across all examples. In particular, we expanded each target $\mathbf{T}_i$ into a matrix of shape

$$896 \times \left( \sum_{\ell=1}^{70} 2d_\ell + 102 \right), \tag{S1}$$

whose first $2d_1$ columns specify the target data for individual 1, next $2d_2$ columns for individual 2, and so forth. The final 102 columns were reserved as *communal* tracks, each corresponding to one of the $2 \times 51 = 102$ possible track types (*coverage* and *junction* data for 51 tissues) across our dataset. For each track in the example, we also populated the communal track of the matching type. Hence, the communal

[5] Sum-aggregation was used for junction data due to its sparsity.

tracks are always (partially) populated regardless of the individual that the example is obtained from, so fitting a model on these tracks requires learning an individual-agnostic sequence-to-phenotype mapping. Finally, a binary mask **M** of the same shape was introduced to indicate which entries in the padded target matrix contain valid data for training. In summary, each dataset example contains two haplotype sequences, their corresponding (padded) target matrices, and a binary mask.

##### 1.2.2 RNA binding proteins

In-vivo binding activity for 150 RNA binding proteins (RBPs) in two cell lines (`K562` and `HepG2`), as measured by enhanced CLIP (eCLIP) assays, was reported in [10]. We downloaded the processed datasets (BED format) from ENCODE [11]. The following 150 RBPs were included:

```
AARS, AATF, ABCF1, AGGF1, AKAP1, AKAP8L, APOBEC3C, AQR, BCCIP, BCLAF1,
BUD13, CDC40, CPEB4, CPSF6, CSTF2, CSTF2T, DDX21, DDX24, DDX3X, DDX42,
DDX51, DDX52, DDX55, DDX59, DDX6, DGCR8, DHX30, DKC1, DROSHA, EFTUD2,
EIF3D, EIF3G, EIF3H, EIF4G2, EWSR1, EXOSC5, FAM120A, FASTKD2, FKBP4,
FMR1, FTO, FUBP3, FUS, FXR1, FXR2, G3BP1, GEMIN5, GNL3, GPKOW, GRSF1,
GRWD1, GTF2F1, HLTF, HNRNPA1, HNRNPC, HNRNPK, HNRNPL, HNRNPM, HNRNPU,
HNRNPUL1, IGF2BP1, IGF2BP2, IGF2BP3, ILF3, KHDRBS1, KHSRP, LARP4, LARP7,
LIN28B, LSM11, MATR3, METAP2, MTPAP, NCBP2, NIP7, NIPBL, NKRF, NOL12,
NOLC1, NONO, NPM1, NSUN2, PABPC4, PABPN1, PCBP1, PCBP2, PHF6, POLR2G,
PPIG, PPIL4, PRPF4, PRPF8, PTBP1, PUM1, PUM2, PUS1, QKI, RBFOX2, RBM15,
RBM22, RBM5, RPS11, RPS3, SAFB, SAFB2, SBDS, SDAD1, SERBP1, SF3A3,
SF3B1, SF3B4, SFPQ, SLBP, SLTM, SMNDC1, SND1, SRSF1, SRSF7, SRSF9,
SSB, STAU2, SUB1, SUGP2, SUPV3L1, TAF15, TARDBP, TBRG4, TIA1, TIAL1,
TRA2A, TROVE2, U2AF1, U2AF2, UCHL5, UPF1, UTP18, UTP3, WDR3, WDR43,
WRN, XPO5, XRCC6, XRN2, YBX3, YWHAG, ZC3H11A, ZC3H8, ZNF622, ZNF800,
ZRANB2.
```

##### 1.2.3 microRNA

The microRNA (miRNA) dataset was generated by curating publicly available AGO2 CLIP-Seq datasets from 12 different cell lines: 22RV1, A2780, A673, BC-1, BC-3, BCBL, DG-75, HCT-116, HEK293, HeLa, hESC, and MCF7 [12–21]. The protocols used were PAR-CLIP, HITS-CLIP, eCLIP and miR-eCLIP [22–25]. The mentioned protocols use an antibody to pull down the AGO2 protein and sequence the corresponding RNA fragments. miR-eCLIP has an additional step where only chimeric reads are kept and both the RNA fragment and the corresponding miRNA are sequenced.

In order to process each dataset, we first trimmed the adapters from the sequenced reads using `fastp` [26] with:

```
fastp --adapter_sequence [...] -w 16
--trim_poly_x --cut_tail 30 --trim_front1 [...] --trim_tail1 0.
```

`adapter_sequence` was set to the corresponding adapter sequence used in each CLIP dataset, and `trim_font1` was set to 5 for PAR-CLIP and 0 for the other protocols. We then aligned the reads to the corresponding reference genome. For PAR-CLIP, we used `bowtie` [27] with:

```
bowtie -v 2 -m 3 --best --strata --threads 32 -S -q.
```

For HITS-CLIP and eCLIP we used `bowtie2` [28] with:

```
bowtie2 -D 20 -R 3 -N 0 -L 20 -i S,1,0.50 -k 4 -p 32.
```

After aligning the reads, we used a peak caller to binarize the read pileups and identify regions of the reference genome that have sufficient read pileups to represent an AGO2 binding site. For PAR-CLIP we used `PARalyzer` [29] peak caller and set the cutoff to a minimum of 5 reads for a region to be classified as a peak. For HITS-CLIP and eCLIP we first deduplicated the reads using `picard` [30] and used the `clipper` [31] peak caller. All the data processing for miR-eCLIP data like adapter trimming, aligning, and peak calling were done by `eclipsebio` [32].

Next, we combined peaks from multiple replicates of the same cell line by taking a union of all the peaks, which we'll refer to as the *target*. For PAR-CLIP there are certain regions of the genome where there is read alignment but not sufficient enough to pass the peak cutoff, these regions were treated as neither negative nor positive. For HITS-CLIP, eCLIP and miR-eCLIP were peaks are then filtered based on the log-p-value generated by the peak caller. Only peaks with $-\log p > 3$ were considered as high confidence while the other peaks were treated as low quality. For this, we created a binary track which will be referred to as the *mask*. For model training, we multiplied the *target* with the *mask*, which effectively set the target value at the masked positions to 0.

#### 1.3 Training and fine-tuning

We trained a seven-model ensemble on the RNA-seq dataset (Section 1.2.1), and then finetuned the first model of the ensemble on the RBP and miRNA datasets (Sections 1.2.2 and 1.2.3). In all subsequent sections, the former will be referred to as the BigRNA ensemble (or simply as BigRNA), and the latter as the fine-tuned model. All training was done on v3-32 TPUs with a batch size of 1 per device, which corresponds to an effective global batch size of 32. We consider an epoch to be 1000 training steps, which corresponds to 32,000 training examples.

**Loss.** We used the masked Poisson loss between the sum of the two predictions and the average of the two targets. Formally, for a fixed example, let $\mathbf{Y}_0$ and $\mathbf{Y}_1$ be the model predictions on both haplotypes, and $\mathbf{T}_0$ and $\mathbf{T}_1$ be the respective targets, where $\mathbf{Y}_i$ and $\mathbf{T}_i$ are of the shape in Supplementary Equation S1. Then the entry-wise losses are computed by:

$$\mathbf{M} \odot \left( (\mathbf{Y}_0 + \mathbf{Y}_1) - \frac{1}{2}(\mathbf{T}_0 + \mathbf{T}_1) \odot \log(\mathbf{Y}_0 + \mathbf{Y}_1) \right), \tag{S2}$$

where $\odot$ denotes an entry-wise Hadamard product and $\log(\cdot)$ is applied entry-wise.

**RNA-seq training.** We used the default Rectified Adam [33] optimizer from TensorFlow Add-ons, a linear learning rate warm-up over 5 epochs, and gradient norm clipping. Since the RNA-seq dataset was not strand-specific, we also applied data augmentation on each example whereby, with probability 0.5, the input DNA sequence was replaced with its reverse-complement and the targets were reversed along the sequence dimension.

Empirically, we found improved performance by ensembling over seven iterations of our model, which used the same base architecture but differ minorly in their choices of training hyperparameters. Supplementary Table S1 summarizes the differences between the models and hyperparameters in our ensemble. We'll refer to this ensemble as $\mathcal{M}$ throughout subsequent sections.

**RBP and miRNA fine-tuning.** We used the default Adam [34] optimizer from TensorFlow, gradient norm clipping at 0.2, and the same random reverse-compliment applied to the RNA-seq dataset. We also applied a random ±64bp jitter to the input sequence. For efficiency, this was done as a preprocessing transform on the dataset: we applied the random shift over the base dataset 10 times independently, and concatenated the results into a 10× larger dataset. The final head layer of the model was first trained for 10 epochs with a learning rate of $2 \times 10^{-4}$, and then the entire model was further trained for another 30 epochs with a learning rate of $10^{-4}$.

#### 1.4 Inference

BigRNA takes as input fixed-length DNA sequences and produces fixed-length outputs at 128bp resolution (Section 1.1). To make predictions on an arbitrary-length genomic interval (<114688bp), we pad it to 196608bp by expanding upstream 40960bp and downstream by the remaining length. The number 40960 comes from the fact that the BigRNA model makes predictions for the center 114688bp subsequence of the input interval, so the outer $\frac{1}{2}(196608 - 114688) = 40960$bp of the input sequence serve as context (Section 1.1). Hence, we are left-aligning the interval of interest to the output of the model. Next, we extract the genomic sequence of the expanded 196608bp interval and pass it through the model to get a $896 \times d$ prediction matrix. Each value in the prediction matrix gets repeated 128 times along the sequence dimension, and the matrix is cropped to be of same length and aligned with the original interval of interest, to facilitate downstream processing. Supplementary Fig. S1 summarizes the inference procedure.

Afterwards, we may apply post-processing. For the ensemble models trained on RNA-seq data, we often subset the predictions to its entries that correspond to some output heads of interest, such as the coverage heads (i.e., all output heads that predict coverage data), junction heads, or coverage or junction heads corresponding to a specific tissue. We always used the heads corresponding to specific individuals rather than the communal heads. In all subsequent sections, we will refer to selection of output heads as $\mathcal{T}$, which corresponds to particular indices in **Supplementary Data 1**:

- Output head selection $\mathcal{T}$: "coverage head" or "coverage data"
  `sample_type == "coverage" AND`
  `averaged_across_multiple_subjects == False`
- Output head selection $\mathcal{T}$: "junction head" or "junction data"
  `sample_type == "junction" AND`
  `averaged_across_multiple_subjects == False`

- Output head selection $\mathcal{T}$: "coverage head in tissue X" or "coverage data in tissue X"
  `sample_type == "coverage" AND`
  `tissue == X AND`
  `averaged_across_multiple_subjects == False`
- Output head selection $\mathcal{T}$: "junction head in tissue X" or "junction data in tissue X"
  `sample_type == "junction" AND`
  `tissue == X AND`
  `averaged_across_multiple_subjects == False`

We can further combine multiple predictions in order to achieve strand-invariance or higher resolutions. We describe these algorithms in the following subsections.

##### 1.4.1 Strand-invariant predictions

The reverse-compliment augmentation during training (Section 1.3) encourages the model to be strand-agnostic. However, to produce a truly strand-invariant prediction, we can average the predictions on both the original interval and the corresponding one on the opposite strand, whose sequence is just the reverse-complement of the original sequence. We always apply strand-averaging unless specified otherwise.

##### 1.4.2 Higher-resolution predictions

We can artificially produce more granular predictions by combining multiple coarser predictions. To predict at an $r$-bp resolution (where $1 \leq r < 128$), we construct $\lceil \frac{128}{r} \rceil$ shifted intervals with step size $r$ and make a 128bp resolution prediction on each one. These predictions are aligned back to the original interval and then min-aggregated, as shown in Supplementary Fig. S2. We use min-aggregation instead of average to avoid over-smoothing, which is important for predicting sharply peaked data such as the RBP and miRNA tracks. Unless specified otherwise, we always predicted at a higher resolution $r = 64$bp.

#### 1.5 Saliency scoring

To guide the design of steric blocking oligonucleotides (SBO) (Section 5), we consider a gradient-based saliency method inspired by [35, 36]. Formally, given a one-hot encoded sequence $\mathbf{X} \in \{0,1\}^{196608 \times 4}$ and BigRNA model $\varphi$, let

$$f(\mathbf{X}) = \texttt{masked\_mean}(\varphi(\mathbf{X})) \in \mathbb{R} \tag{S3}$$

be obtained by passing $\mathbf{X}$ through $\varphi$ and then averaging the subset of output heads of interest. Then, we compute a vector of scores for each token in $\mathbf{X}$ by:

$$\mathbf{G} = \nabla f(\mathbf{X} + \mathbf{X} \odot 0.1\mathbf{E}), \tag{S4}$$

$$\mathbf{s}_{\varphi,\mathbf{X}} = \sum_{i=1}^{4} \mathbf{X}_{\cdot,i} \odot \left( \mathbf{G}_{\cdot,i} - \frac{1}{4} \sum_{k=1}^{4} \mathbf{G}_{\cdot,k} \right) \in \mathbb{R}^{196608}, \tag{S5}$$

where $\mathbf{G}_{\cdot,i}$ denotes the $i$-th column of the matrix $\mathbf{G}$, and $\mathbf{E}$ is a random matrix whose elements are drawn independently from the standard Gaussian distribution.

To score an arbitrary-length interval ($<$114688bp), we can pad it to 196608bp and then obtain position-wise scores as described above. However, this process may be sensitive to how we choose the context surrounding the interval, so to construct a more robust score, we average the scores from multiple such choices. Specifically, we expand the interval such that it begins at the $(40960 + 1 + k)$-th position in $\mathbf{X}$, for 100 equally-spaced values of $k$ ranging from 0 to 2000, inclusive.

### 2 Held-out performance evaluation

To evaluate the performance of BigRNA, we used Gencode v29 [5] to select a held-out set of genes $\mathcal{G}$ that are completely outside of the training and validation set, and each overlaps an interval in the test set.

#### 2.1 Gene expression

This section describes the methodological details relevant to **Fig. 1b-c**.

For a held-out gene and tissue of interest, we predicted the coverage by averaging over all exons $\mathcal{X}$ in the gene, models in the BigRNA ensemble $\mathcal{M}$ (Section 1.3), and coverage output heads $\mathcal{T}$ corresponding to the tissue. Let $\mathbf{y}_{\chi,\tau,\varphi}$ be the output of the head $\tau \in \mathcal{T}$ in the BigRNA model $\varphi \in \mathcal{M}$ on the exon $\chi \in \mathcal{X}$, using the inference procedure from Section 1.4. Then, the predicted coverage is computed as:

$$s = \mu\left(\{\mu\left(\mathbf{y}_{\chi,\tau,\varphi}\right) \mid \chi \in \mathcal{X}, \tau \in \mathcal{T}, \varphi \in \mathcal{M}\}\right), \tag{S6}$$

Here, $\mu(\cdot)$ denotes the mean of entries of its input, and will be used to simplify notation throughout this text: for a finite set $\mathcal{A}$ of scalars or vectors,

$$\mu(\mathcal{A}) = \frac{1}{|\mathcal{A}|} \sum_{\alpha \in \mathcal{A}} \alpha \tag{S7}$$

and for a vector or matrix $\mathbf{A}$, we computed $\mu(\mathbf{A})$ as if $\mathbf{A}$ were a set of its (scalar) entries. The target value is computed similarly but using the processed RNA-seq data instead of model outputs.

#### 2.2 Differential gene expression between tissues

This section describes the methodological details relevant to **Fig. 1e-f**.

To assess performance on predicting differential expression between tissues, we constructed all pairwise tissue comparisons. Given a pair of tissues, let $\mathbf{s}_i \in \mathbb{R}^{|\mathcal{G}|}$ be the predicted coverage across the held-out genes for the $i$-th tissue, which is computed following Section 2.1. Let $\mathbf{t}_i \in \mathbb{R}^{|\mathcal{G}|}$ be the corresponding target coverage, with each $t$ (for one gene) summarized from the RNA-seq dataset in a similar way as the prediction. Then we considered the correlation between the predicted and target log fold changes, $\log_2(\mathbf{s}_1) - \log_2(\mathbf{s}_2)$ and $\log_2(\mathbf{t}_1) - \log_2(\mathbf{t}_2)$, respectively, where $\log_2(\cdot)$ is applied entry-wise.

#### 2.3 *SLC7A8* prediction visualization

This section describes the methodological details relevant to **Fig. 1d**.

We used the GRCh38 genome build and Gencode v29 [5] annotation to query the sequence in the genomic region of *SLC7A8* gene. We made predictions using BigRNA as discussed in Section 1.4. We averaged coverage output heads for "Brain - Hypothalamus" tissue to get a single value per position. For visualization, we multiplied the predicted values by a factor of 2 to represent the predicted total coverage over two haplotypes (reflecting the reference RNA-seq data).

#### 2.4 RNA binding proteins

This section describes the methodological details relevant to **Fig. 1g**.

Using the RBP dataset (Section 1.2.2), we created one binary label for each 128bp window. Over a single window, let $\mathbf{t} \in \mathbb{R}^{128}$ be the targets and $\mathbf{r} \in \mathbb{R}^{128}$ be the average of two low-confidence replicates. We define the binary label as follows:

$$\ell = \begin{cases} 1, & \text{if } \mu(\mathbf{t}) > 0.25, \\ 0, & \text{if } \mu(\mathbf{t}) = \mu(\mathbf{r}) = 0, \\ \text{undefined}, & \text{otherwise}, \end{cases} \tag{S8}$$

where $\mu(\cdot)$ denotes a mean and is defined in Supplementary Equation S7.

We computed the BigRNA predictions as described in Section 1.4 using the fine-tuned model. For DeepRiPe [37], we downloaded the models from their repository[6] and generated predictions for each base by feeding as input to the model a 128bp sequence centered at the base (63bp upstream and 64bp downstream). For all predictions, we took the mean within 128bp window as the predicted value. To compute average precision score for each RBP dataset, we used the output head corresponds to the particular RBP for BigRNA (**Supplementary Data 2**). We were not able to obtain output RBP names from DeepRiPe, so we took the output index that maximize performance of DeepRiPe for each RBP.

#### 2.5 microRNA

This section describes the methodological details relevant to **Fig. 1h**.

We used the 3' UTR sequences of the held-out genes for evaluation. Using the miRNA dataset (Section 1.2.3), we created one binary label for each 128bp window. Over a single window, let $\mathbf{t} \in \mathbb{R}^{128}$ be the targets and $\mathbf{m} \in \{0, 1\}^{128}$ be the mask. We define the binary label as follows:

$$\ell = \begin{cases} 1, & \text{if } \mu(\mathbf{t}) \geq 0.25, \\ 0, & \text{if } \mu(\mathbf{t}) = \mu(\mathbf{m}) = 0, \\ \text{undefined}, & \text{otherwise}, \end{cases} \tag{S9}$$

where $\mu(\cdot)$ denotes a mean and is defined in Supplementary Equation S7.

We computed the BigRNA predictions as described in Section 1.4 using the fine-tuned model. For TargetScan [38], we downloaded the pre-computed predictions from their website.[7] For all predictions, we took the mean within 128bp window

[6] https://github.com/ohlerlab/DeepRiPe/tree/master/Results/Encode_models
[7] https://www.targetscan.org/vert_80/vert_80_data_download/All_Target_Locations.hg19.bed.zip

as the predicted value, for the output head that correspond to the target cell line (**Supplementary Data 2**).

### 3 Variant effect on gene expression

#### 3.1 Pathogenicity evaluation

This section describes the methodological details relevant to **Fig. 2a,d-f**.

##### 3.1.1 Data processing

Single nucleotide variants (SNVs) classified as pathogenic or likely pathogenic (P/LP) in the 3' and 5' UTR were obtained from [39]. The putative benign set of variants were constructed separately for the 3' and 5' UTR. For each transcript with a P/LP variant in its respective UTR, SNVs classified as benign or likely benign in the same UTR of the transcript were obtained from ClinVar (last accessed Oct. 20, 2022). To further supplement the putative benign set, SNVs were obtained from gnomAD v3.0 [40]. For each transcript with a variant classified as P/LP in the corresponding UTR, the genomic coordinates of all exons, based on NCBI RefSeq v110 annotations for *Homo sapiens* [41] within the respective UTR were used to construct an SQL query to extract variants within these regions from `bigquery-public-data.gnomAD.v3_genomes__chr*` tables on BigQuery. Variants were further filtered to have a total allele frequency greater than 0.001 (0.1%). We did this to attempt to match at least one putative benign variant for every UTR, but in some cases this was not possible. This resulted in the final dataset of 20 P/LP and 224 putative benign variants in the 3' UTR and 58 P/LP variants and 120 variants in the 5' UTR (**Supplementary Data 3**).

##### 3.1.2 Predictive scores

**BigRNA.** We used the GRCh38 human reference genome sequence to build an interval centered on the transcript. In the event that the length of the transcript exceeds the model's output length 114688, the interval is defined from the end of the relevant UTR of the dataset with 2,000bp additional genomic sequence beyond the UTR end, and extended into the transcript until the output length 114688 is reached.

We made predictions using BigRNA following Section 1.4 on all coding exon intervals $\mathcal{X}$ within the transcript. We use the ensemble $\mathcal{M}$ (Section 1.3) and only its coverage output heads $\mathcal{T}$. Given a model $\varphi \in \mathcal{M}$, head type $\tau \in \mathcal{T}$, and exon $\chi \in \mathcal{X}$, let $\mathbf{y}_{\chi,\tau} \in \mathbb{R}^L$ be the prediction for head $\tau$ obtained by passing through model $\varphi$ the wild-type exon $\chi$, where $L$ is the length of $\chi$. Define $\mathbf{y}'_{\chi,\tau} \in \mathbb{R}^L$ analogously for the mutant-type (variant) exon. Then the per-model variant effect score is computed as:

$$\bar{\mathbf{y}}_\chi = \mu(\{\mathbf{y}_{\chi,\tau} \mid \tau \in \mathcal{T}\}) \in \mathbb{R}^L, \quad \text{(similarly for } \bar{\mathbf{y}}'_\chi \text{ with } \mathbf{y}'_{\chi,\tau}\text{)}$$
$$s_\varphi = \frac{\max\operatorname{abs}\{\max\operatorname{abs}(\bar{\mathbf{y}}'_\chi - \bar{\mathbf{y}}_\chi) \mid \chi \in \mathcal{X}\}}{\max\{\max(\bar{\mathbf{y}}_\chi) \mid \chi \in \mathcal{X}\} + 10}, \tag{S10}$$

where $\mu(\cdot)$ denotes a mean and is defined in Supplementary Equation S7, and a pseudo-count of 10 is added in the denominator of $s_\varphi$. Here, $\max\operatorname{abs}(\cdot)$ denotes the element with the maximum magnitude with signs preserved; formally, $A = \max\operatorname{abs}(\mathcal{A})$ for a set $\mathcal{A}$ if and only if $A \in \mathcal{A}$ and $|A| \geq |\alpha|$ for all $\alpha \in \mathcal{A}$. If $\mathbf{a}$ is a vector, then $\max\operatorname{abs}(\mathbf{a})$

and max($\mathbf{a}$) are computed as if $\mathbf{a}$ were a set of its entries. Finally, the overall variant effect score is computed by averaging the per-model scores over the ensemble $\mathcal{M}$.

**Enformer.** Enformer weights were loaded from [2]. For each variant, two sequences encompassing Enformer's full context length were constructed: one centered at the reference allele and one centered at the alternative allele. For each sequence, predictions for the forward and reverse strand were made by Enformer, and the average was taken.

We made use of predictions from a subset of output heads $\mathcal{T}$ that correspond to CAGE data within a local window of size 640 centered around the variant. For a head type $\tau \in \mathcal{T}$, let $\mathbf{y}_\tau \in \mathbb{R}^{640}$ be the prediction for head $\tau$ by passing through the wild-type sequence. Similarly define $\mathbf{y}'_\tau \in \mathbb{R}^{640}$ for the mutant-type (variant) sequence. We compute the variant effect score as:

$$s = \mu(\{\ell(\mathbf{y}'_\tau) - \ell(\mathbf{y}_\tau) \mid \tau \in \mathcal{T}\}), \qquad \ell(\mathbf{y}) = \log_2\left(\sum_{i=1}^{640} \mathbf{y}_i\right) + 1 \in \mathbb{R}, \tag{S11}$$

where $\mu(\cdot)$ denotes a mean and is defined in Supplementary Equation S7, and 1 is added as a pseudo-count. This mirrors what was done by [2, 42] for variant effect predictions.

**Saluki.** For each 3' UTR variant, we constructed a 6-dimensional track for the Saluki [43] model, consisting of the one-hot encoded DNA sequence of the transcript, the coding frame, and the splice site positions for the wild-type (reference) and mutant-type (alternative) sequence, as described by the authors. We used 50 cross-fold validation models (which we will refer to as the ensemble $\mathcal{M}_s$) provided by the authors. For each model in the ensemble $\varphi \in \mathcal{M}_s$, let $y_\varphi \in \mathbb{R}$ be the prediction from passing the wild-type track through the model. Define $y'_\varphi \in \mathbb{R}$ similarly for mutant-type. The variant effect score $s$ is computed as the difference between the two, mean-aggregated over the ensemble:

$$s = \mu(\{y'_\varphi - y_\varphi \mid \varphi \in \mathcal{M}_s\}), \tag{S12}$$

where $\mu(\cdot)$ denotes a mean and is defined in Supplementary Equation S7.

**FramePoolCombined.** Predictions for 5' UTR variants were made using the Kipoi interface provided for the FramePoolCombined model [44, 45] using the GRCh38 reference FASTA(last accessed Aug. 1, 2023) and GTF file of the GRCh38 NCBI RefSeq (last accessed Aug. 1, 2023) table was downloaded from UCSC. Each variant was predicted one at a time in its own VCF to yield individual variant effects. The predicted mean ribosome load fold change reported by the model was used as the variant effect score.

##### 3.1.3 Evaluation - Statistical analysis

The receiver operator characteristic (ROC) curve and precision-recall curve (PRC) was plotted for each predictor on each dataset and the area under the curves (AUC) were calculated. Each curve was bootstrapped 10,000 times and the standard deviation from the bootstrap results was calculated. The mean true positive rate (TPR) with one standard deviation and the mean precision with one standard deviation is plotted. To calculate the significance between AUROC and AUPRC differences of the models, a permutation test using 10,000 permutations was performed with the $p$-value being reported.

##### 3.1.4 Evaluation - Variants of unknown significance (VUS) analysis

Variants from ClinVar (last accessed Apr. 30, 2023) that were classified as VUS and in 3' and 5' of the UTRs of the transcripts that they were reported in were extracted. These variants were further filtered to those that were in the same genes as those included in the benchmark datasets. These variants were predicted and scored with BigRNA in the same manner as described in Section 3.1.2. Scores for these variants are provided in **Supplementary Data 4**.

#### 3.2 *NAA10* case example

This section describes the methodological details relevant to **Fig. 2b**.

We made predictions using the BigRNA model (Section 1.4) on both wild-type and mutant-type (variant) sequences around the *NAA10* gene sequence. We took the average across all coverage outputs, resulting in one track for the wild-type prediction and one track for the variant prediction for visualization.

#### 3.3 Predicting the impact of disrupting polyadenylation sites

This section describes the methodological details relevant to **Fig. 2c**.

To evaluate BigRNA's ability to predict polyadenylation (poly(A)) sites, we conducted an *in-silico* 11bp (a symmetric 5bp expansion centered on a 1bp site) N-mask tiling analysis across each poly(A) region. Poly(A) sites were obtained from PolyASite 2.0, a curated database containing inferred poly(A) sites from publicly available 3' end sequencing datasets [46]. PolyASite 2.0 was further filtered to include only those from human, located in terminal exons, and belonging to transcripts that were less than 114,000bp to ensure that the entire transcript would be within BigRNA's output length. From this filtered dataset, we selected the top 200 poly(A) sites with the highest average expression across all samples reported in PolyASite 2.0.

To define the poly(A) regions, we expanded each poly(A) signal sequence from PolyASite 2.0 by ±100bp to cover proximal regulatory elements. Subsequently, we generated 11bp N-masked tiles across each resulting 206bp region, treating each tile like a variant.[8] For each tile, we calculated the predicted change in gene expression by comparing the BigRNA predictions for the N-mask (variant) and the poly(A) signal sequence (wild-type).

We made predictions using BigRNA as described in Section 1.4, on the transcript plus 500bp downstream.[9] We used the first model in the ensemble and all output heads $\mathcal{T}$. Given a head type $\tau \in \mathcal{T}$, let $\mathbf{y}_\tau \in \mathbb{R}^L$ be the prediction for head $\tau \in \mathcal{T}$ obtained

[8] This effectively sets the one-hot encoded input to all zeros within the N-masked region.
[9] To capture expression/isoform changes that might occur if there is a shift to a more distal polyA isoform.

by passing through the wild-type transcript, where $L$ is the transcript length. Similarly define $\mathbf{y}'_\tau \in \mathbb{R}^L$ for the mutant-type (variant) transcript. We computed one score for each variant by mean-aggregating over the transcript length and all output heads:

$$s = \mu\left(\left\{\mu\left(\mathbf{y}_\tau\right) - \mu\left(\mathbf{y}'_\tau\right) \mid \tau \in \mathcal{T}\right\}\right), \tag{S13}$$

where $\mu(\cdot)$ denotes a mean and is defined in Supplementary Equation S7. This yields one track for each selected gene. For visualizing the average prediction across all selected genes (**Fig. 2c**, bottom), we normalize the scores by their maximum magnitude across all genes.

To visualize changes in expression for all possible point mutations around the polyadenylation site (PAS) of *NAA10* (**Fig. 2c**, top), we used *NAA10* PAS and its surrounding 100bp on either end and performed saturation mutagenesis by point-mutating every nucleic acid base to every other nucleic acid base. The scoring was done in the same way as mentioned above.

#### 3.4 Expression quantitative trait loci (eQTL) evaluation

This section describes the methodological details relevant to **Fig. 2g**.

##### 3.4.1 Data processing

To evaluate the ability of BigRNA to predict distal regulatory variation, we created a benchmark of fine-mapped expression quantitative trait locus (eQTL) variants from GTEx v8 [3]. This dataset contains true positive (TP) examples which consist of eQTLs and true negative (TN) examples which have been matched for impactful characteristics [47]. Publicly available fine-mapped eQTLs from the eQTL Catalog [48] were used to define the positive class, which used SuSiE [47] to derive credible sets from cis eQTL (±1 megabase (Mbp) of eGene) association statistics.[10] To arrive at a set of high-confidence putative causal eQTL, we restricted the gene-variant pairs to associations with a posterior inclusion probability (PIP) greater than 0.95 in at least one tissue, nominally associated with gene expression ($p < 5 \times 10^{-6}$), and amenable to BigRNA prediction (±50kbp of transcription start site of the corresponding eGene). The true negative class consisted of variants that were not associated ($p > 0.05$) with expression of cis (±1Mbp) genes in any tissue from GTEx and amenable to BigRNA prediction (±50kbp of transcription start site for the APPRIS [49] principal isoform of the eGene). To avoid class imbalance and ensure similar characteristics between the two groups, each true positive was matched to a true negative based on minor allele frequencies (±10%) and distance to transcription start site (±10kbp) in the same gene (Supplementary Fig. S11). The benchmark dataset consisted of 1,374 unique variants in the true positive class and 1,162 unique variants in the true negative classes with a PIP greater than 0.5. The benchmark contained 397 unique variants in the true positive class with a PIP of greater than 0.95 and 356 paired negative control variants. We used the higher confidence PIP threshold of 0.95 for subsequent analyses as this indicates

[10] eQTL Catalog SuSIE Fine-mapping: http://ftp.ebi.ac.uk/pub/databases/spot/eQTL/susie/

statistical confidence that each variant used in the positive set is causally driving the association signal up to the definition of the $\rho$-credible set with here $\rho$ set to 0.95.

We defined distal and proximal eQTLs based on standard nomenclature with a 10 kilobase (Kbp) cut off [50–52]. We calculated the distance between the variant position in GRCh38 and the transcription start site of the eGene (or the eGene of the paired positive eQTL variant). We used the APPRIS transcript of the eGene[11] for transcription start site calculations in all cases. This procedure led to 409 proximal and 344 distal examples for bench-marking within subsets.

###### 3.4.2 Predictive scores

We used BigRNA to predict the impacts of true positive and negative eQTL variants on the expression of their eGenes. To do so, we used the GRCh38 human reference genome sequence and the Gencode v41 [5] annotation set[12] to build an interval of length 114688bp (equal to the output length of BigRNA) centered on the variant. We built intervals for Enformer in a similar way.

Using a similar strategy as [2], we computed the scores as described in Section 3.1.2 and Supplementary Equation S10, matching tissue type to the tissue where the positive control has been fine-mapped as causally impacted expression of the eGene (**Supplementary Data 5**). For more details on other scoring methods we considered, see Scoring considerations for eQTL evaluation.

###### 3.4.3 Evaluation

To calculate precision-recall (PRC) and receiver-operator (ROC) curves we used the scores calculated in the previous section to segregate positive and negative, where labels corresponding to the status of the variant as a fine-mapped eQTL or a negative control.

[11] APPRIS Transcript Definitions: https://appris.bioinfo.cnio.es/

[12] Gencode v41 annotation set: https://www.gencodegenes.org/human/release_41.html

### 4 Variant effect on splicing

#### 4.1 Intron retention

This section describes the methodological details relevant to **Fig. 3d**.

##### 4.1.1 Data processing

Splicing variants [53] were downloaded from the SPCards database (release v1.0).[13] Intron retention variants were manually curated by searching for entries with evidence for intron retention, such as "...retains upstream intron...". Each entry was individually curated to ensure that the source publication supports the reported effects. Partial intron retention events (e.g. extension of the upstream/downstream exon partially into the intron) were excluded. For each variant in the above dataset, we searched within 50bp local neighbourhood to create a matching negative set of variants that do not cause intron retention using gnomAD [54] v2.1. To deplete the negative set of variants that may cause deleterious effects, we require the allele frequency to be larger than 0.001 in any population. After combing all positive and negative variants, we filtered out the trivial variants that overlap the core dinucleotide.[14]

##### 4.1.2 Predictive scores

**BigRNA.** For each variant in the dataset, we made predictions using BigRNA (Section 1.4) on both wild-type and mutant-type sequences around the affected intron. We used the ensemble $\mathcal{M}$ (Section 1.3) and only its coverage output heads $\mathcal{T}$. Given a head type $\tau \in \mathcal{T}$ and model $\varphi \in \mathcal{M}$, let $\mathbf{y}_i$, $\mathbf{y}_u$, and $\mathbf{y}_d$ be the predictions for $\tau$ obtained by passing to $\varphi$ the wild-type intron, upstream exon, and downstream exon, respectively. First, we compute a ratio between the intron and its two flanking exons:

$$r_{\tau,\varphi} = \frac{\text{median}(\mathbf{y}_i)}{\frac{1}{2}(\max(\mathbf{y}_u) + \max(\mathbf{y}_d)) + 1}, \tag{S14}$$

where median and max are taken over the entries of its argument. Define $r'_{\tau,\varphi}$ similarly using predictions on the mutant-type sequences. The score is computed by aggregating over output heads and models in ensemble by:

$$s = 1 - \min_{\varphi \in \mathcal{M}} \mu\left(\left\{ \frac{r_{\tau,\varphi}}{r'_{\tau,\varphi}} \,\middle|\, \tau \in \mathcal{T} \right\}\right), \tag{S15}$$

where $\mu(\cdot)$ denotes a mean and is defined in Supplementary Equation S7.

**SpliceAI.** We used the five-model ensemble from [55].[15] Given a SpliceAI model $\varphi$, for each variant, we get the acceptor and donor strength prediction for both wild-type and mutant-type by predicting on sequence centered at the affected intron. Let the

[13] http://www.genemed.tech/spcards/download
[14] Core dinucleotide refers to GT on the donor site and AG on the acceptor site.
[15] https://github.com/Illumina/SpliceAI

predicted acceptor and donor strength for wild-type sequence be $p_a$ and $p_d$. Similarly for mutant-type we have predictions $p'_a$ and $p'_d$. We compute the difference between wild-type and mutant-type predictions:

$$s_\varphi = \frac{p_a + p_d}{2} - \frac{p'_a + p'_d}{2}. \tag{S16}$$

The score is then computed as a mean-aggregated over the models $\varphi$ in ensemble.

Curated variants, with target labels and scores can be found in **Supplementary Data 8**.

##### 4.1.3 Evaluation

We used ROC curves to evaluate model performance on classifying positive and negative variants. To construct a confidence interval around each ROC curve, we boostrapped 1000 times and used the 34.1-th and 84.1-th percentiles as the lower and upper bounds, which correspond to $\pm 1$ standard deviation from the mean in a Gaussian distribution.

#### 4.2 Exon skipping - MaPSy evaluation

This section describes the methodological details relevant to **Fig. 3a**.

##### 4.2.1 Data processing

The MaPSy dataset [56] was downloaded from the MaPSy challenge website.[16] Exonic disease mutations were screened in mini-gene system, both in-vivo (transfection in tissue culture) and in-vitro (incubation in cell nuclear extract). The challenge is to predict the degree to which a given variant causes changes in splicing. For each mutation, MaPSy reports four values from the in-vivo assay: input wild-type $a$, spliced wild-type $b$, input mutant $c$, and spliced mutant $d$. We compute the splicing odds ratio $R$ with $\chi^2$ 95% confidence intervals $[C^{\ell}_{\text{vivo}}, C^{u}_{\text{vivo}}]$:

$$\begin{aligned} R &= \frac{(a+\varepsilon)(d+\varepsilon)}{(b+\varepsilon)(c+\varepsilon)}, \\ \sigma &= \sqrt{\frac{1}{a+\varepsilon} + \frac{1}{b+\varepsilon} + \frac{1}{c+\varepsilon} + \frac{1}{d+\varepsilon}}, \\ C^{\ell}_{\text{vivo}} &= \exp\left(\log R - 1.96\sigma\right), \\ C^{u}_{\text{vivo}} &= \exp\left(\log R + 1.96\sigma\right), \end{aligned} \tag{S17}$$

where $\varepsilon = 0.1$ is a pseudocount for numerical stability. For the same mutation, MaPSy also reports a quartet from the in-vitro assay, on which we can again use Supplementary Equation S17 to compute an analogous confidence interval $[C^{\ell}_{\text{vitro}}, C^{u}_{\text{vitro}}]$.

To label each mutation as skipping or non-skipping, we used the above confidence intervals from both in-vivo and in-vitro assays. Given a specified skipping level $p$, the

[16] http://www.genomeinterpretation.org/cagi5-mapsy.html

mutation was assigned the binary label:

$$\ell_p = \begin{cases} \text{skipping}, & \text{if } C^u_{\text{vivo}} < p \text{ and } C^u_{\text{vitro}} < 1, \\ \text{non-skipping}, & \text{if } p \leq C^\ell_\nu < 1 < C^u_\nu \leq \frac{1}{p} \text{ for } \nu \in \{\text{vivo}, \text{vitro}\}, \\ \text{undefined}, & \text{otherwise.} \end{cases} \tag{S18}$$

For example, if $p = 0.5$ (50% skipping), to label the mutation as $\ell_{0.5} = \text{skipping}$, we required that the in-vivo upper confidence limit to be less than 0.5, while applying a less stringent threshold of 1 on the in-vitro upper confidence limit. To label the mutation as $\ell_{0.5} = \text{non-skipping}$, we required both confidence intervals to be within $(0.5, 2)$. To filter out mutations with low confidence, we generated binary labels at multiple skipping levels $p \in \{0.5, 0.35, 0.3, 0.25, 0.2, 0.1\}$ and filtered out mutations that were labelled undefined at *all* levels. The processed high confidence dataset consists of 1816 mutations. For each of these mutations, we reconstructed the genomic coordinates of their affected exon by aligning the artificial construct sequence to the reference genome and locating the donor and acceptor sites.

##### 4.2.2 Predictive scores

**BigRNA.** For each mutation in the dataset, we made predictions using BigRNA (Section 1.4) on both the wild-type and mutant-type sequences on the affected exon. We used the ensemble $\mathcal{M}$ (Section 1.3) and only its junction output heads $\mathcal{T}$. Given a head type $\tau \in \mathcal{T}$ and BigRNA model $\varphi \in \mathcal{M}$, let $\mathbf{y}$ be the predictions for the head obtained by passing the wild-type sequence through $\varphi$, and define $\mathbf{y}'$ similarly for the mutant-type. Then we compute the score as:

$$s_{\tau,\varphi} = \max\left(\frac{\mathbf{y}}{\mu(\mathbf{y})} - \frac{\mathbf{y}'}{\mu(\mathbf{y}')}\right), \tag{S19}$$

where $\mu(\cdot)$ denotes a mean and is defined in Supplementary Equation S7. We computed the overall score by averaging over the collection of output heads $\mathcal{T}$ and models $\mathcal{M}$ in the ensemble.

**Enformer [2].** We used all CAGE output heads, computed the per-head score according to Supplementary Equation S19, and averaged them over output heads to get a scalar score.

**SpliceAI [55].** We downloaded the command line tool from their repository.[17] For each mutation, the tool reports four positive values: acceptor gain `ds_ag`, acceptor loss `ds_al`, donor gain `ds_dg`, and donor loss `ds_dl`. We first added back the sign for the score corresponding to "gain": −`ds_ag`, `ds_al`, −`ds_dg`, `ds_dl`, and took the score with the maximum absolute value[18].

[17] https://github.com/Illumina/SpliceAI
[18] max abs as defined in Supplementary Equation S10.

##### 4.2.3 Evaluation

A comparison of the three models for classifying skipping versus non-skipping variants is shown in Supplementary Fig. S7, in which we show the other two models in addition to BigRNA (**Fig. 3a**). Target value is the skipping binary label at threshold 0.5, as defined in Section 4.2.1, and predictive scores are computed as described in Section 4.2.2 for each of the 3 models. Permutation test and $p$-value on ROC difference was computed as described in Section 3.1.3, with $100,000$ permutations. Curated variants with all predictive scores can be found in **Supplementary Data 7**.

#### 4.3 *ACADM* variant effect on TDP-43 binding

This section describes the methodological details relevant to **Fig. 3b**.

We made predictions using the fine-tuned BigRNA (Section 1.3), on both the wild-type and the mutant-type (variant) sequence containing the c.468+7A>G variant on the *ACADM* gene (Section 1.4). We visualized the output corresponding to the TDP-43 binding profile in K562 cells for each of the wild-type and variant sequence predictions (**Fig. 3b, top**). We then used the BigRNA model trained on RNA-seq (Section 1.3) to predict on the same wild-type and variant sequences. We used the first model in ensemble $\mathcal{M}$ (Section 1.3) and only its coverage output heads $\mathcal{T}$. Predictions were averaged across output heads to yield an one-dimensional track for visualization (**Fig. 3b, bottom**).

#### 4.4 Validating variant effect of *ATP7B* mutation

This section describes the methodological details relevant to **Fig. 3c**.

We used CRISPR-Cas9 editing to generate HepG2 hepatoblastoma derived lines carrying the *ATP7B* `c.3243+5G>A` mutation. We designed a CRISPR RNA (crRNA) to cut at chromosome 13 position 52518240 (GRCh37). We designed 99bp single stranded oligodeoxynucleotides (ssODNs) containing the mutation of interest as the template for knock-in via homology directed repair. We conducted nucleofection using the Amaxa 4D nucleofector to introduce the crRNA and ssODNs to wild-type HepG2 cells. After the cells recovered from the transfection, we performed limited dilution by diluting the pool of edited cell lines to 5 cells/well and seeding into a 96 well (96W) plate. We isolated genomic DNA and sent samples for Sanger sequencing to determine genotype. We used a compound heterozygous line with a large deletion in the exon body which has ∼20% wild-type *ATP7B*.

For confirmation of exon 14 skipping, total RNA was extracted from the homozygous *ATP7B* `c.3243+5G>A` HepG2 cell line 48 hours after seeding. cDNA was produced using first-strand synthesis (High-Capacity cDNA Reverse Transcription Kit; Life Technologies) and used as a template to perform PCR with primers targeting exons 13 and 15 (Fw: CACGGCTGTCATGGTGGG, Rev: CTGACTTTGCACCCAATTCC). PCR products were analyzed by agarose gel electrophoresis.

We made predictions using BigRNA (Section 1.4) on both the wild-type and mutant-type sequences on the affected exon. We used the ensemble $\mathcal{M}$ (Section 1.3) and only its coverage output heads $\mathcal{T}$. Predictions were averaged across output heads and models in the ensemble to yield a one-dimensional track for visualization.

#### 4.5 Validating variant effect of *ABCA4* mutation

This section describes the methodological details relevant to **Fig. 3e**.

WERI-Rb-1 cells containing the *ABCA4* `c.5714+5G>A` mutation were generated using CRISPR-Cas9 gene editing. A crRNA was designed to cut at chromosome 1 position 94476351 (GRCh37). 100bp ssODNs containing the mutation of interest were designed as the template for knock-in via homology directed repair. 200,000 WERI-Rb-1 cells (ATCC) were nucleofected with the cRNA and ssODNs using an Amaxa 4D nucleofector and incubated for 72 hours at 37 °C, 5% CO2. Following incubation, cells were transferred to 12 well plates. Cells were harvested after 18 days and plated at 1 or 5 cells per well in 96-well plates. After 30 days, clonal populations were harvested. gDNA was extracted from the cells using the Geneaid Tissue/Blood DNA Mini Kit and long-range PCR (Fw: CCTCTTCTCCCTGCAGTTCG, Rev: GGTCAGGGAGTCAAACCAGG and Fw: TCTGACTCCACTGACAGAGG, Rev: TGGAGGGATGACCATAGAGC) was performed to verify the edit. Samples were run on the MinION (Oxford Nanopore Technologies) to verify that a homozygous edit was generated. For RNA-seq, RNA was extracted from cells using the RNeasy Mini Kit (Qiagen). RNA-seq libraries were prepared with the NEBNext Ultra II Directional RNA preparation kit, with polyA selection performed using the NEBNext Poly(A) mRNA Magnetic Isolation module. Sequencing was carried out on a NovaSeq 6000 sequencer. Reads were aligned with HISAT2 v2.1.0 [7] and per-base coverage was calculated using the `genomecov` command in bedtools v2.30.0 [57]. Coverage of the replicates was averaged ($n = 3$).

We made predictions using BigRNA (Section 1.4) on both the wild-type and mutant-type sequences on the affected exon. We used the ensemble $\mathcal{M}$ (Section 1.3) and only its coverage output heads $\mathcal{T}$. Predictions were averaged across output heads and models in the ensemble to yield a one-dimensional track for visualization.

### 5 Designing steric blocking oligonucleotides (SBO)

#### 5.1 Scoring SBOs for splicing effects

This section describes the methodological details relevant to **Fig. 4a**.

**Predicting an SBO as a variant.** To predict the effect of each SBO, we simulate it as a substitution variant. We replace genomic sequences within the SBO hybridization region with a stretch of N's of the same length as the region. This effectively sets the one-hot encoded input to all zeros within the SBO hybridization region.

**BigRNA predictive score.** For each SBO in the dataset, we simulated it as a variant as mentioned above. BigRNA scores were computed according to Section 4.2.2.

#### 5.2 LabChip exon skipping

This section describes the methodological details relevant to **Fig. 4b**.

All target genes were screened in the HEK293T cell line (ATCC). Cells were seeded at a density of 50,000 or 300,000 cells and transfected with 50 or 300pmol of SBO, respectively. Total RNA was isolated 48h post-transfection and RT-PCR was performed using high-capacity cDNA kit (Promega) and oligo-dT (IDT). PCR was performed with custom primers designed to amplify around the exon of interest. The PCR product was analyzed on an automated capillary electrophoresis separation (Perkin Elmer LabChip-GX). The Labchip Software was used to extract high-resolution sizing and quantitation for all DNA fragments, and a proportion spliced in (PSI) value was calculated for each exon. SBOs were scored using BigRNA as described in Section 5.1.

**SpliceAI predictive score.** For each SBO in the dataset, we simulated it as a variant as mentioned in Section 5.1. We made prediction using SpliceAI [55] model by passing the sequence centered at the affected exon. Given a SpliceAI model $\varphi$, let $p_a$ be the predicted probability at the acceptor site of the wild-type sequence and $p_d$ be the prediction at the donor site. Define $p'_a$ and $p'_d$ analogously for the mutant-type. Then, we compute the score as:

$$s_\varphi = \max(\ell(p_a) - \ell(p'_a), \ell(p_d) - \ell(p'_d)), \tag{S20}$$

where $\ell(p) = \log((p+\varepsilon)/(1-p+\varepsilon))$ is the logit, or inverse-sigmoid function, and $\varepsilon = 10^{-10}$ is added for numerical stability. We further averaged the scores across a five-model ensemble to obtain the final score for the SBO. The models were obtained from the SpliceAI repository.[19]

For both models, Spearman correlation was calculated between the predicted SBO effect and the average PSI-value.

#### 5.3 Screening *ATP7B* exon 6 M645R SBOs

This section describes the methodological details relevant to **Fig. 4c**.

[19] https://github.com/Illumina/SpliceAI. We were not able to directly apply the SpliceAI VCF tool, since it does not work with variants where the reference or alternate allele are longer than 1bp.

In order to measure the effect of SBOs designed to rescue the splicing deficits caused by the `c.1934T>G` mutation in *ATP7B*, compound heterozygous HepG2 cells were generated. This monoclonal cell model harbored a knockout of one copy of *ATP7B* via a large insertion of sequence in exon 6[58], and the 'Spanish' variant `c.1934T>G` on the remaining copy.

Using these cells, a library of 55 SBOs were screened using quantitative PCR (qPCR). Designed SBOs were reverse transfected into the mutant HepG2 cells using Lipofectamine™ RNAiMAX. Following transfection, the cells were incubated for 48 hours, after which, cells were lysed in RLT buffer and RNA was extracted via the RNeasy kit using the Qiacube automated system. RNA concentrations were determined using an Agilent bioanalyzer (RNA nano). First-strand synthesis was performed using a high capacity cDNA kit following the manufacturer's recommendations.

To measure the effect of the SBOs, a qPCR assay was developed to measure "full length" *ATP7B* transcripts which included exons 5, 6, and 7 (Fw: ATTGAG-GAAATTGGCTTTCATGC, Rev: ACAGGAAAGACTTCTTCCACTGC). *ATB7B* expression was normalized to the expression of the housekeeping gene *TBP* (Fw: GC-CCGAAACGCCGAATATA, Rev: CGTGGCTCTCTTATCCTCATGA). qPCR was performed using a 2× SYBR mix with the designed primers. Conditions were based on the Quantstudio 5 qPCR machine.

SBOs were scored using BigRNA as described in Section 5.1. Spearman correlation was computed between the BigRNA prediction and the average qPCR fold change. For visualization purposes, BigRNA predictions were re-scaled to have a minimum of zero and an 80th percentile matching the observed average fold change values.

#### 5.4 Visualization of BigRNA predictions for wildtype vs. M645R variant vs. oligonucleotide and M645R variant in *ATP7B*

This section describes the methodological details relevant to **Fig. 4d**.

For the M645R[20] variant, we generated three sequences encompassing BigRNA's full context length: one centered at the reference allele (wild-type), one centered at the alternative allele (variant), and one centered at the alternative allele with the sequence corresponding to the designed hybridization site of the SBO (variant with SBO), as described in Section 5.1. For each sequence, predictions using BigRNA were generated as described in Section 1.4. We used the ensemble $\mathcal{M}$ (Section 1.3) and only its coverage output heads $\mathcal{T}$. Predictions were averaged across output heads and models in the ensemble to yield one-dimensional tracks (one for wild-type, one for variant, and one for variant with SBO) for visualization.

In addition to visualizing BigRNA's prediction, we also assessed different model's FPR at which this variant can be discovered as an exon skipping variant. We computed BigRNA, SpliceAI and Enformer scores for the M645R variant in *ATP7B*, as discussed in Section 4.2.2. BigRNA reports a score of 0.1586, which corresponds to discovery at 0.7% FPR (Supplementary Fig. S7). For SpliceAI, the score is 0.22 and this variant can be discovered at 0.6% FPR. The Enformer score is 0.0045 and this variant can be discovered at 21.02% FPR.

[20] M645R is `chr13:52535985:A>C` (h37) or `chr13:51961849:A>C` (h38).

#### 5.5 Identification of "N=1" variants associated with autosomal recessive disorders

This section describes the methodological details relevant to **Fig. 4e-f**.

To generate a set of genes which are associated exclusively with autosomal recessive (AR) disorders, we filtered OMIM [59] (last accessed Apr. 19, 2023) to select for AR and exclude other inheritance modes. This resulted in a set of 2563 genes. Next, we collected all pathogenic or likely pathogenic variants in the AR genes from ClinVar (last accessed Apr. 30, 2023; $n = 101,448$). GnomAD [40] allele frequencies (v3.1.2) were employed to subset to variants with very low total worldwide disease prevalence ($\rho \leq 50$), using the following estimates:

$$\rho_h = f_a^2 \times P, \tag{S21}$$

$$\rho_c = 2 \times f_a \times f_o \times P, \tag{S22}$$

$$\rho = \rho_h + \rho_c, \tag{S23}$$

where $\rho_h$ is homozygous prevalence, $\rho_c$ is compound heterozygous prevalence, $f_a$ is the total allele frequency for the variant, $f_o$ is the total allele frequency for other pathogenic/likely pathogenic variants in the gene, and $P$ is an estimate for the global population (8 billion).

These 5486 variants from 1549 genes were further filtered to select SNVs with homozygous prevalence equal to 0 and allele frequencies less than or equal to 0.00002 in all individual GnomAD subpopulations, resulting in a final set of 1582 "N=1" pathogenic variants from 863 genes associated with AR disorders. Synonymous variants ($n = 14$), tolerated missense variants (SIFT [60] scores $> 0.05$, $n = 56$), intronic variants (>8bp from splice sites, $n = 15$), and splice region variants ($\leq$8bp from splice sites, excluding the core dinucleotides, $n = 22$) were prioritized for exon skipping SBO design. Variants with annotation and BigRNA scores can be found in **Supplementary Data 9**.

The `c.2481-12A>G` variant in *MYO1E* (`chr15:59163315:T:C` h38) was predicted to cause skipping of *MYO1E* exon 23. To assess whether the variant effect may be amenable to rescue with an SBO, we performed an in-silico design of all possible 20-mers within 200bp of exon 23, as described in Section 5.1.

To visualize prediction of the lead SBO, we generated three sequences encompassing BigRNA's full context length: one centered at the reference allele (wild-type), one centered at the alternative allele (variant), and one centered at the alternative allele with the sequence corresponding to the designed hybridization site of the SBO (variant with SBO), as described in Section 5.1. For each sequence, predictions using BigRNA were generated as described in Section 1.4. We used the ensemble $\mathcal{M}$ (Section 1.3) and only its junction output heads $\mathcal{T}$. Predictions were averaged across output heads and models in the ensemble to yield one-dimensional tracks (one for wild-type, one for variant, and one for variant with SBO) for visualization.

#### 5.6 Scoring expression increase SBOs

This section describes the methodological details relevant to **Fig. 4g-h**.

To evaluate the ability of BigRNA to identify SBOs that increase gene expression, we used a combination of established saliency mapping techniques (Section 1.5), to evaluate the contribution of each base in a transcript to the gene's expression in human liver (by using a subset of outputs that correspond to liver coverage data in Supplementary Equation S3), yielding a sensitivity score for each base's impact on expression levels. Using this score, we ranked relevant inhibitory regions across the transcript of interest by their propensity for SBO-induced gene expression increase. To calculate the inhibitory score for an SBO, the minimum value of any overlapping bases was taken as the overall SBO score. For lead SBOs and hits in *SMN2*, *ATP7B*, *PRRT2*, and *SERPING1*, the inhibitory score was calculated for all candidate SBOs of the same length targeting anywhere in the APPRIS principal transcript. The scores of background SBOs and hits were compared with a Mann-Whitney U-test.

##### 5.6.1 SBO screen for *PON1*

A library of 2981 steric blocking oligonucleotides (SBOs) with phosphorothioate backbone and 2'-O-methoxyethyl (PS-MOE) chemistry were designed against the human *PON1* gene, using a mix of proprietary machine learning models, experimental datasets, and random tiling. The 2981 SBOs were reverse transfected into primary human hepatocytes (PHH) at a dose of 100 nM. 18,000 PHH cells per well were reverse transfected and plated in 384-well Collagen-I plates. PHH cells (BioIVT) were thawed using Cryopreserved Hepatocyte Recovery Medium (Thermofisher) and plated with InVitroGRO CryoPlating Hepatocyte Medium (BioIVT). All SBOs and small interfering RNAs (siRNAs) were reverse transfected using Opti-MEM reduced serum medium (Thermofisher) coupled with Lipofectamine™ RNAiMAX Transfection Reagent (Thermofisher). Subsequent media changes (every two days) for cell maintenance were performed using Cellartis Power Primary HEP Medium (Takarabio) over a culture window of 7 days. After 7 days, cells were lysed with AlphaLysis buffer (Perkin Elmer) supplemented with 1X HALT protease inhibitor with no EDTA (Thermofisher). Lysis was done by one freeze-thaw cycle at $-80$ °C followed by thorough mixing of the lysates. Expression of *PON1* was evaluated by AlphaLISA and expression levels were compared to a non-targeting control SBO (19-mer designed to have no binding sites at edit distance 2 or lower) to generate fold change values.

The 264 SBOs with highest fold change from the initial screen were taken into a hit confirmation screen, following the same protocol as the initial screen. The top 44 SBOs from the hit confirmation, along with one non-targeting control, were selected and further validated in an eight-point dose-response curve from 400 nM to a 3.125 nM concentration.

Dose-response SBOs were identified as a hit if the max effect exceeded 1.3 fold increase in *PON1* expression and the half maximal effective concentration (EC50) was between 10 nM and 100 nM.

##### 5.6.2 SBO screens for *ATP7B*, *PRRT2*, and *SERPING1*

SBOs targeting *PRRT2* and *SERPING1* were screened by transfection at 50, 200, and 500 nM in Kelly and HuH-7 cells, respectively. SBOs targeting *ATP7B* were screened by transfection at 50 and 200 nM in HepG2 cells. Each cell line was edited to insert a HiBiT tag into the protein of interest[61], in order to measure endogenous protein levels using luminescence. HiBiT tagged cells were created using CRISPR-Cas9 gene editing. A crRNA (IDT technologies) was designed to cut near the 5' end of each target gene and a 190bp single-stranded oligonucleotide donor, containing the 33bp HiBiT sequence and ~80bp of homology arm sequence, was designed as the donor template for each target gene. Test and control SBOs, along with siRNAs, were reverse transfected into HiBiT edited cells using the lipofectamine reagent RNAiMAX. 48 hours following transfection, PrestoBlue HS (ThermoFisher) reagent was added to each well, and incubated following the manufacturer's instructions, in order to measure cell viability. Following incubation, a portion of the media/PrestoBlue mixture was removed and pipetted into black 96 well plates. Fluorescence measurements at 560 nm excitation and 590 nm emission were made on either a Biotek Neo 2 or a Glomax Discoverer instrument.

After fluorescence measurements, the remaining media/PrestoBlue reagent was aspirated and the wells were gently washed once with phosphate-buffered saline (PBS). A master mix of the HiBiT lytic buffer+reagents was made (Promega) and supplemented with 1:100 large BiT (18kDa subunit of NanoBiT luciferase) protein and 1:50 of NanoGlo HiBiT lytic substrate. PBS was aspirated from the washed wells, and the supplemented HiBiT lytic buffer was added. Plates were incubated on a rocking platform for 10 minutes and 50 $\mu$L volumes from each well were removed and pipetted into opaque white 96 well plates. Luminescence readings were made using either a Biotek Neo 2 or a Glomax Discoverer instrument. The luminescence reading was then divided by cell viability reading and normalized to non-targeting control SBOs, in order to calculate a fold change for each SBO. SBOs with average fold change above 1.5 across the three doses and average viability greater than 60% of non-targeting controls were classified as hits.

### Supplementary Figures

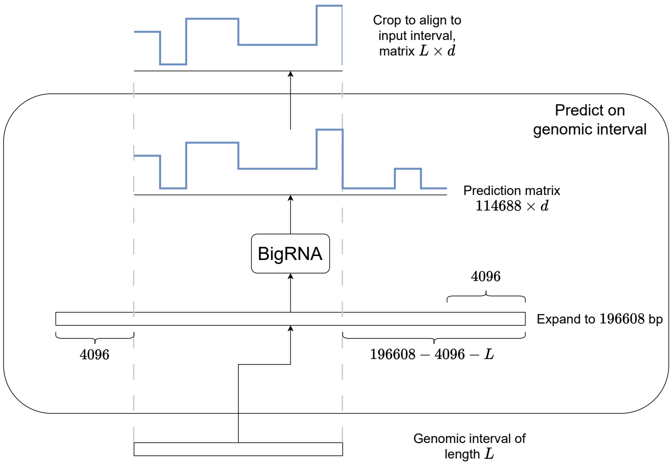

**Supplementary Figure S1** BigRNA prediction on a genomic interval.

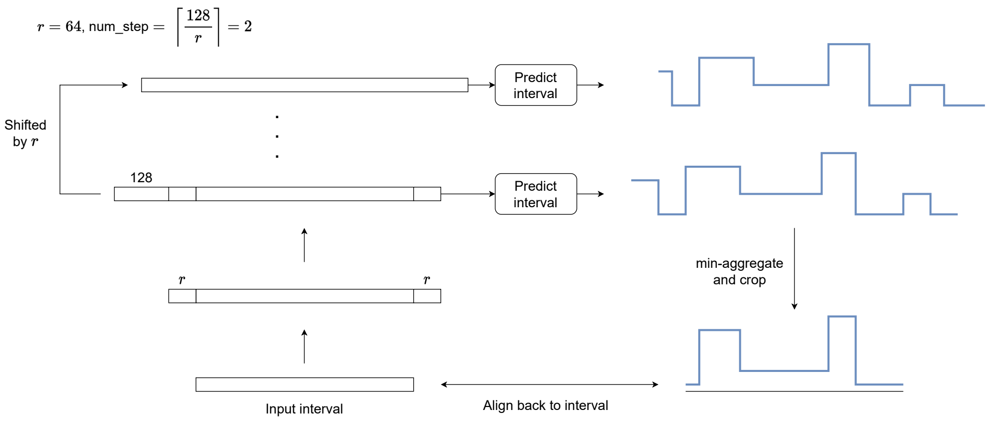

**Supplementary Figure S2** A high-resolution prediction, specifically, with resolution $r = 64$ bp. This extends the procedure for predicting on a genomic interval that is described in Supplementary Fig. S1.

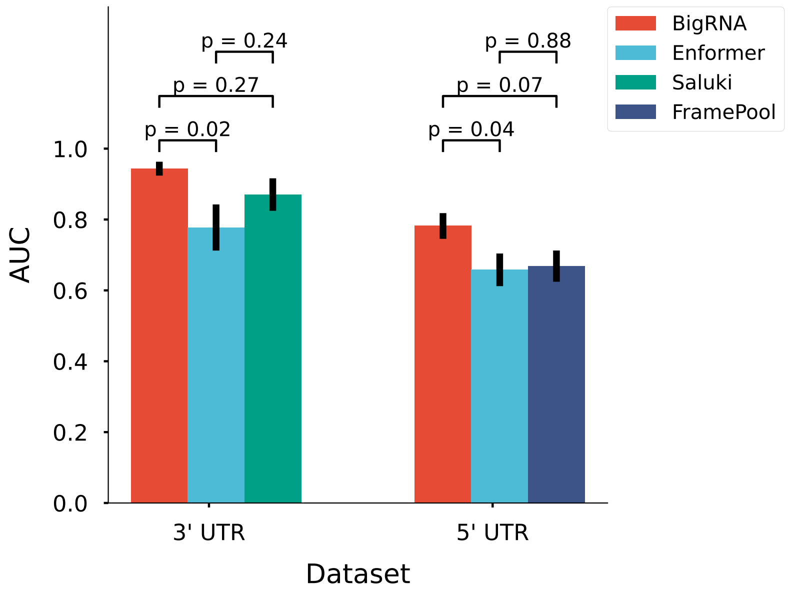

**Supplementary Figure S3** Performance of BigRNA as compared to other deep learning models on the ClinVar P/LP and putative benign variants benchmark. The area under the receiver operator characteristic curve (AUROC) of each of the models at the classification task in the 3' and 5' UTR is plotted. The $p$-value from a permutation test ($n = 10,000$) comparing the AUROC of BigRNA to each of the other models' AUROC is annotated.

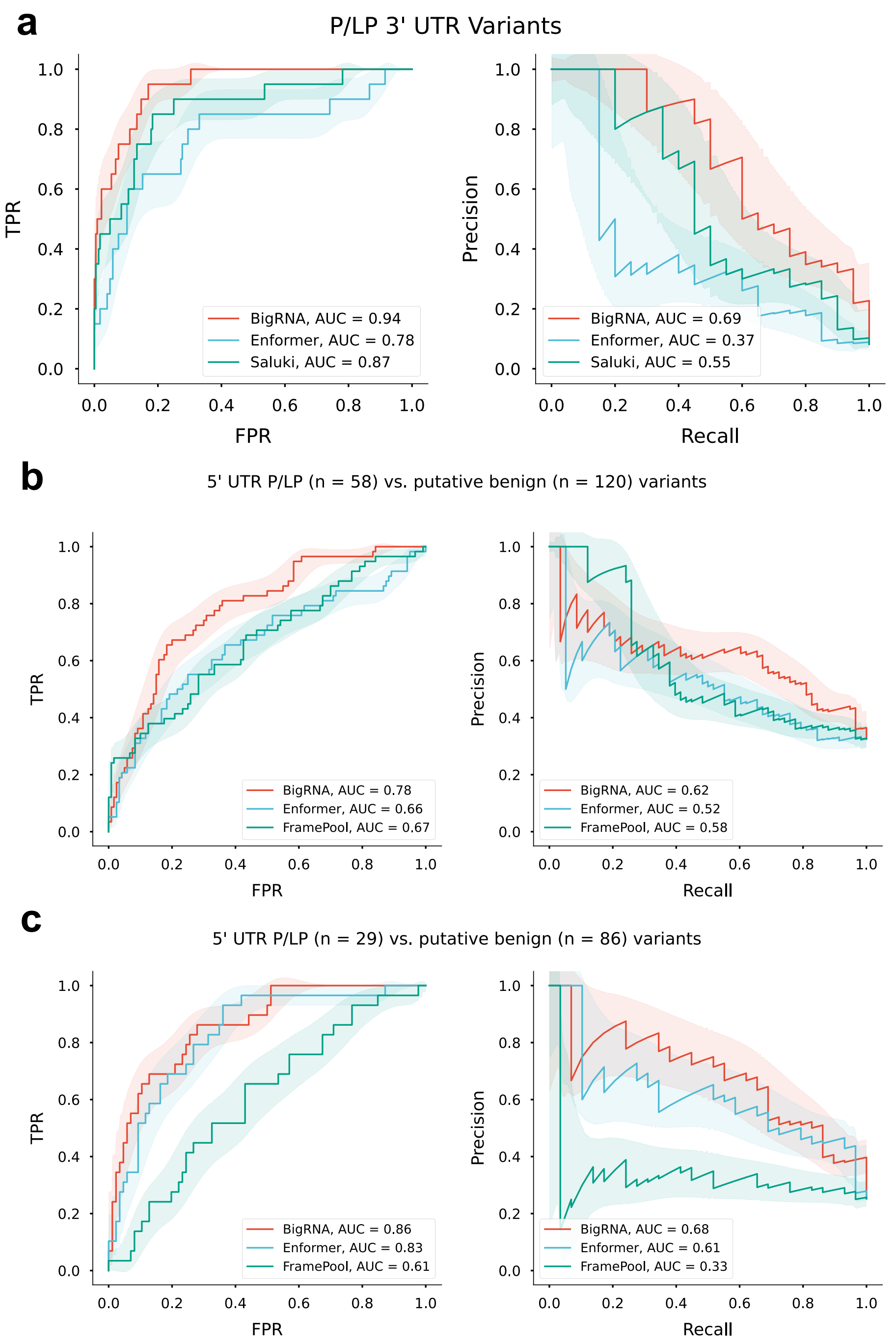

**Supplementary Figure S4** Receiver operator (ROC) and precision-recall (PRC) curves showing the performance of BigRNA and other deep learning models at classifying pathogenic/likely pathogenic (P/LP) variants from ClinVar and putative benign variants in the 3' UTR **(a)**, 5' UTR for all P/LP variants **(b)** and 5' UTR P/LP variants that impact expression **(c)**. **(a)** All P/LP classified SNVs by [39] in the 3' UTR ($n = 20$) against all the matched putative benign variants from our dataset ($n = 224$). BigRNA has a significantly higher AUROC than Enformer ($p = 0.02$, 10,000 permutations) and is comparable to Saluki ($p = 0.27$, 10,000 permutations). **(b)** All P/LP classified SNVs by [39] in the 5' UTR ($n = 58$), regardless of mechanism against all the matched putative benign variants from our dataset ($n = 120$). **(c)** The dataset is subset to just P/LP variants that impact expression (as defined by altering transcription) or those whose mechanism is undefined ($n = 29$) and matched putative benign variants in the matched UTR of this subset ($n = 86$). BigRNA's AUROC is comparable to Enformer ($p = 0.60$, 10,000 permutations) and is significantly higher than FramePoolCombined ($p = 0.002$, 10000 permutations). The fill around the lines represents one standard deviation of the true positive rate and precision, respectively, as calculated from 10,000 bootstraps. AUROC, area under the receiver operator curve; AUPRC, area under the precision-recall curve; TPR, true positive rate; FPR, false positive rate

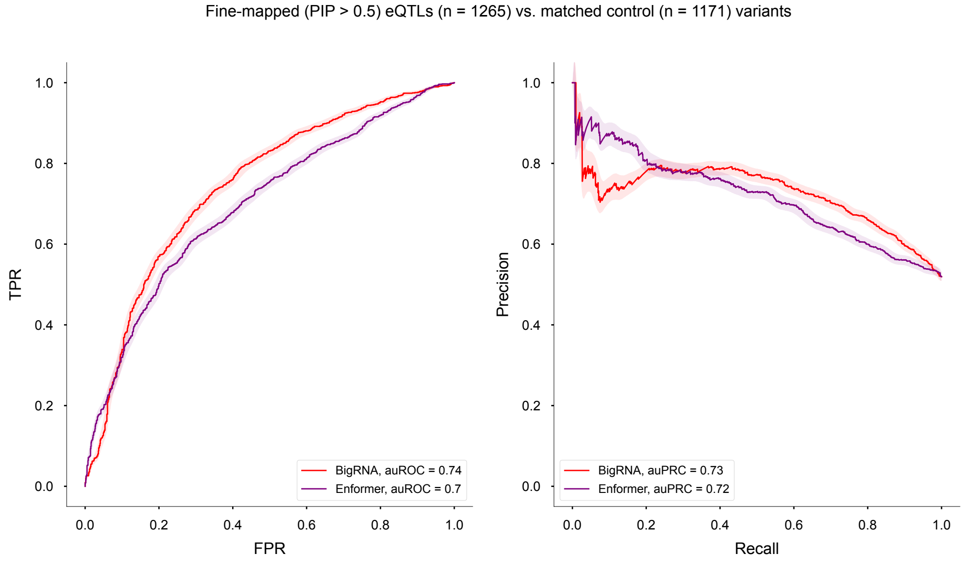

**Supplementary Figure S5** Performance of BigRNA and Enformer on the general task of eQTL classification from matched controls on the entire constructed dataset. We use 10,000 bootstrap resampling iterations to establish significance between the estimates and find that BigRNA significantly out-performs Enformer on the area under the receiver operator characteristic curve (AUROC 0.74 versus 0.70, $p = 0.0047$) however the models are not significantly different on the area under the precision recall curve (AUPRC 0.73 vs 0.72, $p = 0.64$). Both models are displayed using tissue-specific scores derived from the contiguous coding sequence and max aggregation was used over BigRNA ensembles. For more details, see Scoring considerations for eQTL evaluation.

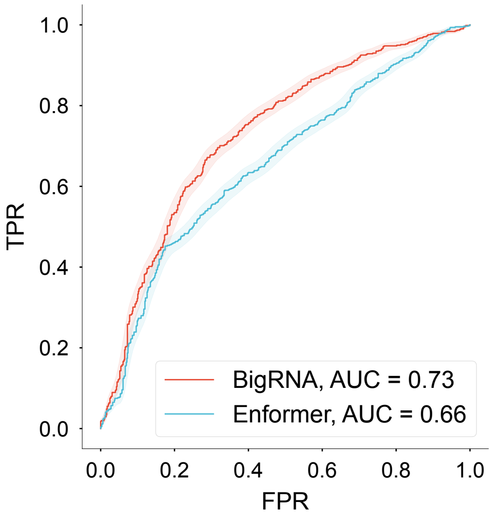

**Supplementary Figure S6** Performance of BigRNA at distinguishing distal (>10 kb) expression quantitative trait loci (eQTLs) loci from matched negative controls compared to Enformer ($p = 10^{-3}$ for difference). eQTLs were scored by using absolute differences in the contiguous coding sequence (CDS) of the effector genes, across all affected tissue outputs, and taking the maximum difference across all BigRNA models in the ensemble.

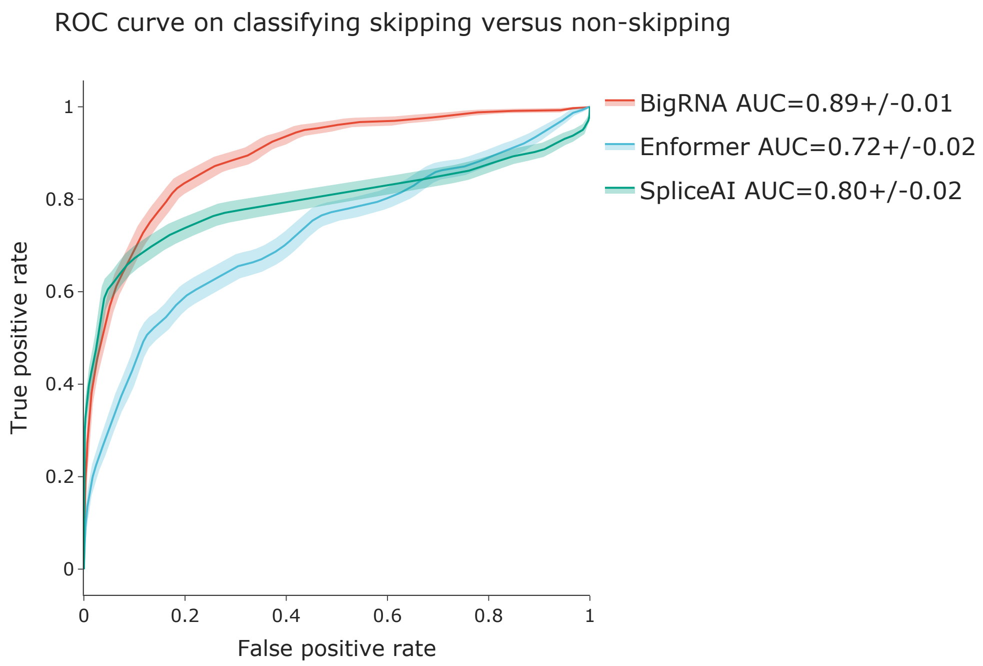

**Supplementary Figure S7** ROC of classifying skipping versus non-skipping variants from the MaPSy dataset at a 50% skipping level.

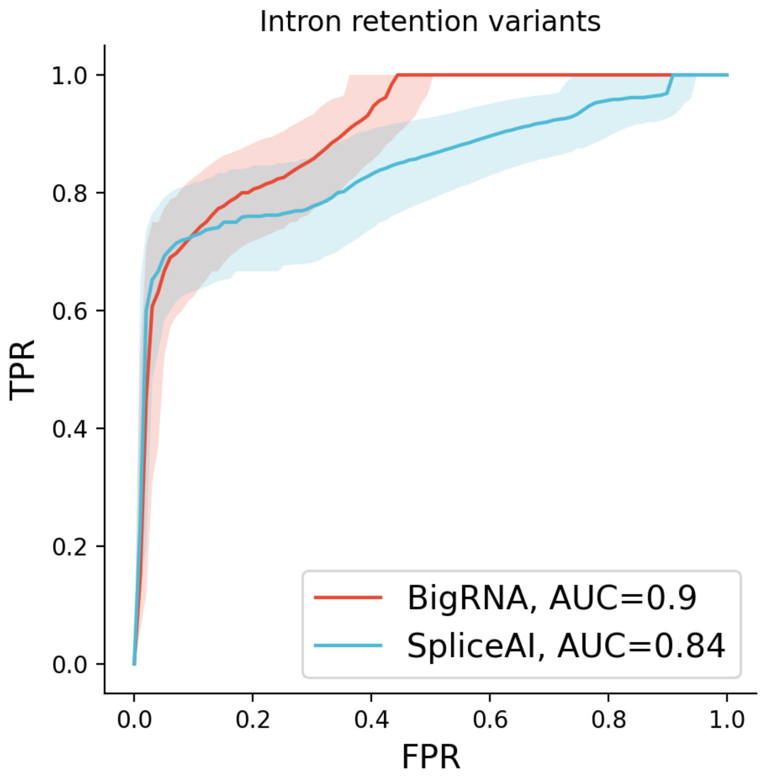

**Supplementary Figure S8** Comparison with SpliceAI for classifying variants that cause intron retention ($n = 25$) from a set of matched variants that do not impact splicing ($n = 63$). The differences do not meet statistical significance.

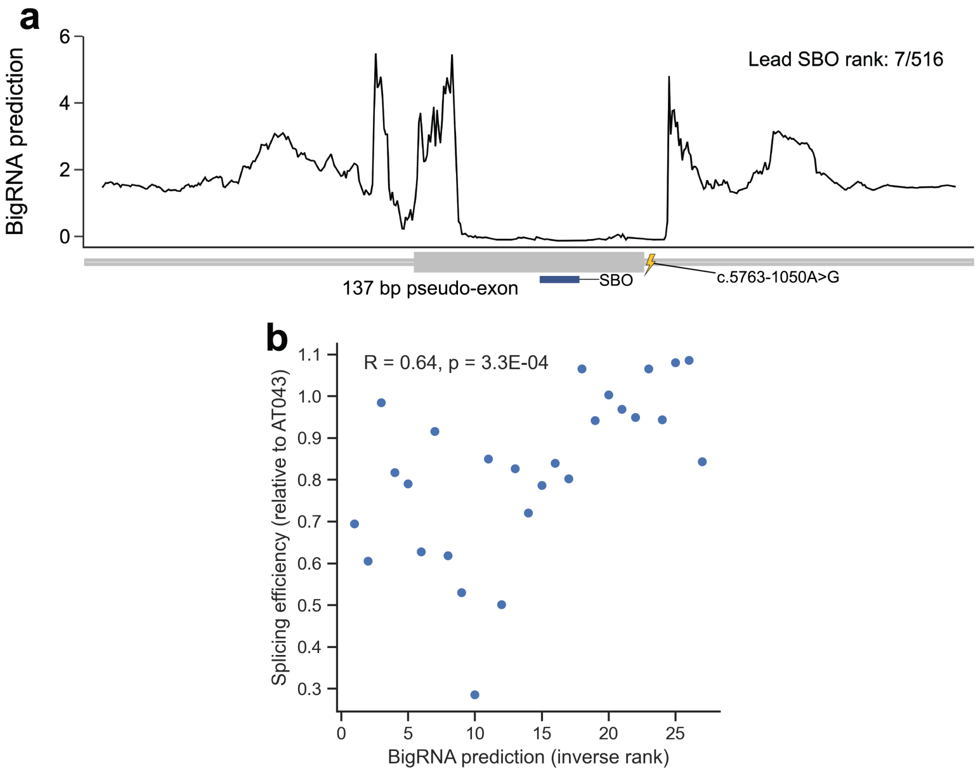

**Supplementary Figure S9** **a.** Predicted effect of SBOs to remedy inclusion of a pseudo-exon in *ATM*, caused by the `c.5763-1050A>G` variant. The blue box shows the position of the lead SBO relative to the pseudo-exon. **b.** Spearman correlation between predictions and experimentally observed splicing efficiencies of 27 SBOs from Kim et al. 2023 [62]. The x-axis shows the relative rank of the BigRNA predictions for the 27 SBOs.

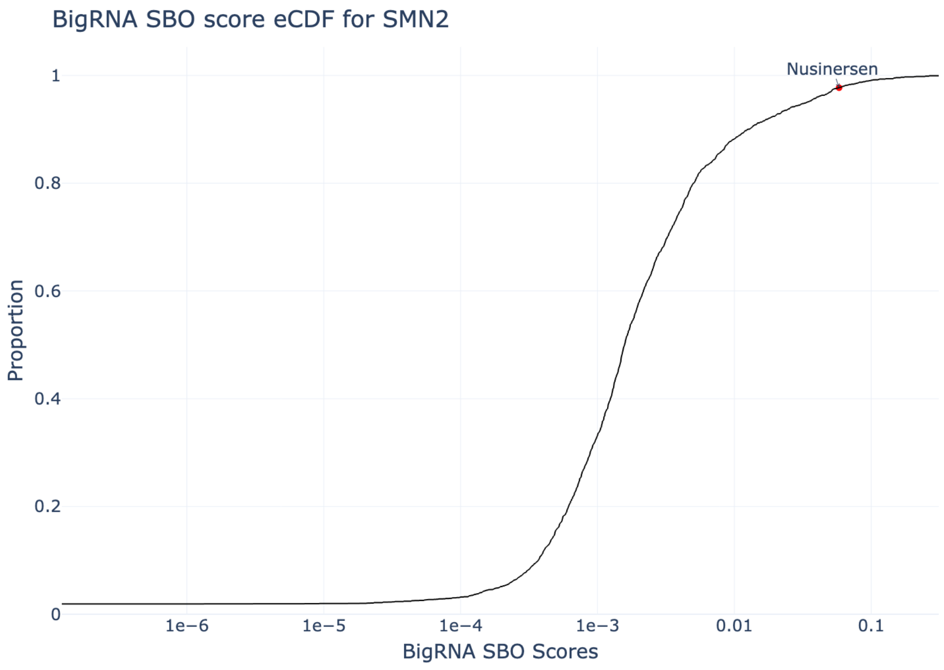

**Supplementary Figure S10** BigRNA inhibitory scores of all 26,901 possible 18-mers targeting *SMN2*. Nusinersen is highlighted with a red dot, ranking in the top 2.28%.

### Supplementary Tables

**Table S1** Training hyperparemters for the seven submodels of our individual-aware ensemble, in ascending order of recency. Notably, model 6 was initialized from the trained Enformer weights [2]. All models are trained for a fixed number of epochs but we only retain the checkpoint with the best validation performance, which is measured at the end of each epoch using the Pearson correlation between the model's predictions and targets on a few validation batches.

| Model No. | 0 | 1 | 2 | 3 | 4 | 5 | 6 |
|---|---|---|---|---|---|---|---|
| Initialization | random | random | random | random | random | random | Enformer |
| Max. epochs | 180 | 170 | 180 | 100 | 100 | 100 | 100 |
| Learning rate | $10^{-4}$ | $5 \times 10^{-5}$ | $5 \times 10^{-5}$ | $10^{-4}$ | $5 \times 10^{-5}$ | $2 \times 10^{-5}$ | $2 \times 10^{-5}$ |
| Grad. Clip | 0.2 | 0.05 | 0.05 | 0.1 | 0.1 | 0.1 | 0.05 |

### Supplementary Notes

#### Scoring considerations for eQTL evaluation

As a part of Section 3.4 a number of considerations were made for the exact scoring methodology from raw predictions. This note describes these considerations and their impacts on the observed performance of both BigRNA and Enformer.

To construct the evaluation benchmark, each positive control variant (fine-mapped eQTL) was matched with a negative control variant with similar minor allele frequency distance to the transcription start site of the effector gene (eGene). This was accomplished by iterating over each of the positive control variants (here defined as having a causal impact on expression of an (eGene) with a PIP $\geq 0.5$ in one or more tissues). The threshold of 0.5 was selected to indicate that the variant was the most probable causal variant in its credible set. Additional fine-tuning downstream was performed to narrow to the set of confident variants with a PIP threshold of $\geq 0.95$, indicating that the variant was suspected of being the sole candidate causal variant in its credible set. Matching was performed in a lenient manner to allow for the maximum number of negative control variants from the same eGene. Minor allele frequencies were set to match within 10% of the positive variant, and distance to TSS was set to match within 5 kilo-basepairs (kbp) (Fig. S11). This matching procedure produces 1211 pairs of positive / negative variants for the same eGene.

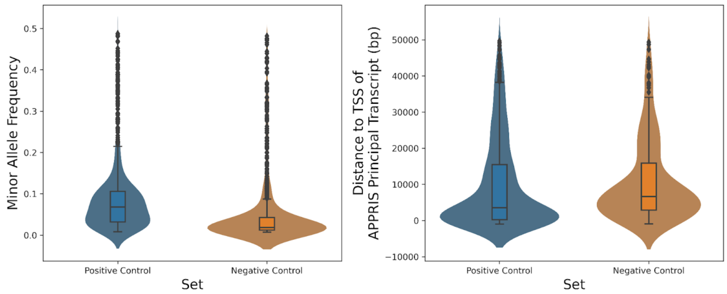

**Supplementary Figure S11** Distribution of positive eQTL variants and matched negative controls for minor allele frequency (MAF) and distance to the transcription start site of their effector gene (eGene).

We use BigRNA and Enformer to predict the effects of these variants. We capture 100kb of sequence prediction centered on the variant of interest, creating an output matrix for scoring $\phi$ with dimensionality Number of variants $\times$ Number of output heads $\times$ Number of base pairs. Denote these dimensions $V$, $H$, and $B$ respectively. To construct a score from $\phi$ which intuitively represents the magnitude of the difference between the reference and the alternate prediction, consider the two output matrices for the reference sequence ($\phi^R$) and the sequencing containing the alternate, effect, allele ($\phi^A$). Each eQTL has one or more "tissue(s) of origin" where it has been fine-mapped as putatively causal. Consider for a particular variant $v \in 1, \ldots, V$ an indicator variable $T_v$ which is an $H$-vector (that is, an indicator variable on the output heads corresponding to the ones which are informative for the tissue of origin of variant $v$). The number of such mapping tissues is available in Fig. S12. For variant $v$ we take the effective matrix of tissue-specific predictions for the alternate allele $\phi^A_{v,h\in T_v}$ and the reference allele $\phi^R_{v,h\in T_v}$ which is of dimensionality $1 \times \sum T_v \times B$. Take $f$ to be an arbitrary aggregation function which operates over the $H$ axis of $\phi^R_{v,h\in T_v}$ and consider two possible definitions (the mean and the max). We wish to use $f$ to aggregate each $\phi_{v,h\in T_v}$ such that it is of dimesionality $1 \times B$. We use the given aggregation function to transform $\phi_{v,h\in T_v}$ to a $1 \times B$ vector for scoring denoted as $\gamma^A$ or $\gamma^R$ for alternate prediction and reference prediction respectively.

$$\gamma_b^{A,R} = \begin{cases} \text{Mean}, & \frac{1}{\sum_i T_v} \sum_{i=0}^{H} \phi^{A,R}_{v,i,b} \\ \text{Max}, & \max_h \phi^{A,R}_{v,h,b} \end{cases}$$

In this formulation we consider then the $B$-vectors $\gamma^A$ and $\gamma^R$ for each variant $v \in V$ for the purposes of scoring. The indices of these aggregated vectors represent base pair positions from the output of each deep learning model. Note that after experimentation, we see little difference in the performance of either BigRNA or Enformer as a consequence of the decision of aggregation function $f$ at this early stage (data not shown). To construct a score for the variant we consider a subset of this vector which is indicative of coding region of the eGene such that $C \subseteq B$. Note that the choice of $C$ as the coding region is arbitrary but was arrived at after empirical testing. An alternate formulation would take $C$ as a single index corresponding to the transcription start site and expand a fixed window around the index as in [2]. Later we empirically evaluate the decision to use the contiguous coding sequence rather than this transcription start site approach. The "score" is then a function of the two aggregated tracks such that

$$\begin{aligned} \gamma^D &= \gamma^A_{b\in C} - \gamma^R_{b\in C} \\ S(\gamma^D, \gamma^R_{b\in C}) &= \frac{\gamma^D_{\arg\max_i |\gamma^D_i|} - \gamma^R_{\arg\max_i |\gamma^R|}}{\gamma^R_{\arg\max_i |\gamma^R|} + c} \end{aligned}$$

where $c$ is a constant. We use $c = 10$ throughout for this evaluation.

An additional area of nuance is that this score must be seperately calculated for each of the seven "ensemble" models which compose BigRNA. This gives us a score $S_v$ for each variant $v$ which is a function of its aggregated reference and alternate $B$-vectors $\gamma^R$ and $\gamma^A$ respectively. We must make a decision on a second aggregation function to combine these seven scores, here called $S_v^e$ for each ensemble model $e \in 0, \ldots, 7$. We experiment with two such methods, namely the mean of the ensemble models and the max of the ensemble models. An interpretation of these two scores would be that the first captures the collective information from all ensemble nodes, while the second indicates whether an effect is indicated in any ensemble node individually. This would then simply be

$$S_v^{\text{mean, max}} = \begin{cases} \text{mean,} & \frac{1}{7}\sum_{i=0}^{7} S_v^i \\ \text{max,} & S_v^{\arg\max_i |S_v^i} \end{cases}$$

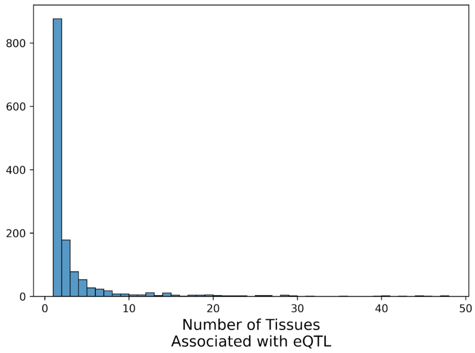

**Supplementary Figure S12** Distribution of the number of effector tissues where each of the positive eQTL variants had been fine-mapped.

We perform permutation bootstrapping to assess the improvement above Enformer for various strategies including $S_v^{\text{mean}}$ versus $S_v^{\text{max}}$ and find that $S_v^{\text{max}}$ leads to the highest classification accuracy for the eQTL classification task (see Fig. 3.4 for more details on evaluations, Supplementary Fig. S2). We assess other parameter's impact on the delta between the two models and additionally find significant differences between the two primarily in the category with more eQTLs (PIP = 0.5) for distal eQTLs (here defined as those which are more than 10 kilobases from their eGene). These classifications do not reflect differences in scoring, but rather simply selecting different sets of $v \subseteq V$.

To motivate the decision to calculate scores specific to a particular set of output heads relevant to the tissue-of-origin ($T_v$ above being an indicator vector for each variant $v$ of relevant heads) we compare the performance of a matched tissue model to an agnostic one (with all entries of $T_v$ set to 1). Note that even when $T_v$ is a 1-vector, we still score using only coverage heads from RNA-seq data (or matched CAGE-seq heads

| PIP | TSS | Score | Δ AUROC | Δ AUROC P | Δ AUPRC | Δ AUPRC P |
|---|---|---|---|---|---|---|
| 0.5 | Distal | Mean | -0.04 | 0.059 | -0.043 | 0.144 |
| 0.5 | Distal | Max | -0.072 | 0.001 | -0.059 | 0.051 |
| 0.5 | Proximal | Mean | 0.003 | 0.87 | 0.033 | 0.193 |
| 0.5 | Proximal | Max | -0.022 | 0.282 | 0.026 | 0.334 |
| 0.5 | All | Mean | -0.019 | 0.185 | 0.0 | 0.992 |
| 0.5 | All | Max | -0.046 | 0.002 | -0.01 | 0.6 |
| 0.8 | Distal | Mean | -0.042 | 0.155 | -0.044 | 0.276 |
| 0.8 | Distal | Max | -0.075 | 0.012 | -0.06 | 0.148 |
| 0.8 | Proximal | Mean | 0.008 | 0.739 | 0.031 | 0.324 |
| 0.8 | Proximal | Max | -0.018 | 0.496 | 0.029 | 0.425 |
| 0.8 | All | Mean | -0.019 | 0.323 | -0.001 | 0.977 |
| 0.8 | All | Max | -0.044 | 0.025 | -0.006 | 0.818 |
| 0.95 | Distal | Mean | -0.047 | 0.215 | -0.05 | 0.341 |
| 0.95 | Distal | Max | -0.061 | 0.129 | -0.051 | 0.355 |
| 0.95 | Proximal | Mean | 0.017 | 0.574 | 0.04 | 0.353 |
| 0.95 | Proximal | Max | -0.01 | 0.782 | 0.034 | 0.461 |
| 0.95 | All | Mean | -0.014 | 0.563 | 0.004 | 0.897 |
| 0.95 | All | Max | -0.033 | 0.203 | 0.002 | 0.959 |

**Table S2** 10,000 permutation tests were used to establish the difference between the area under the receiver operator curve (AUROC) and area under the precision recall curve (AUPRC) for BigRNA scored using one of three posterior inclusion probability (PIP), three distance to TSS, and two ensemble aggregation methods.

from Enformer, see **Supplementary Data 5**). We compare classification performance on the main eQTL dataset (PIP threshold of 0.5) and find a statistically significant difference between tissue-informed and tissue-agnostic scores for both BigRNA and Enformer (P = 0.0027 and P = 0.0026 respectively from 10,000 iterations of bootstrap resampling; Supplementary Fig. S13)

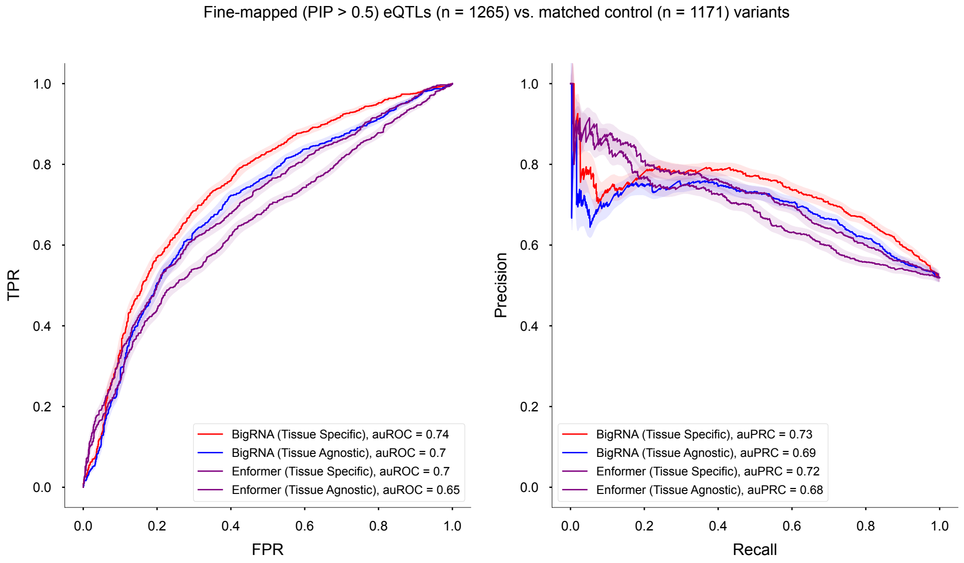

**Supplementary Figure S13** Precision recall and receiver operator characteristic curves for BigRNA and Enformer using tissue-specific and tissue-agnostic scoring methods. We use 10,000 bootstrap resampling iterations to assess significant differences and find that the area under the receiver operator characteristic curve (AUROC) is significantly improved by using tissue-specific rather than tissue agnostic scores for BigRNA ($p = 0.0029$) and Enformer ($p = 0.0015$) but not the area under the precision recall curve (AUPRC) for either BigRNA ($p = 0.056$) or Enformer ($p = 0.071$). For BigRNA, max aggregation over output heads was used along with the CDS (rather than TSS) score.

In addition to the aggregation strategy and tissue-specificity, the choice of $C$ (the region to evaluate for the purposes of scoring) has an impact on the classification performance. To assess which of these scores is empirically supported by the data, we calculate two versions of the score for both BigRNA and Enformer. We use the Appris transcript (if available) for each of the eGenes (or eGene of matched eQTL in the case of negative control variants) to compute a score as previously defined across the contiguous coding sequence (CDS). We additionally take the transcription start site of the eGene and create a window 196bp on either side for score computation (TSS). We compare the performance of these scores and find that in both cases the CDS score out-performs the TSS score for both AUROC and AUPRC (Supplementary Fig. S14).

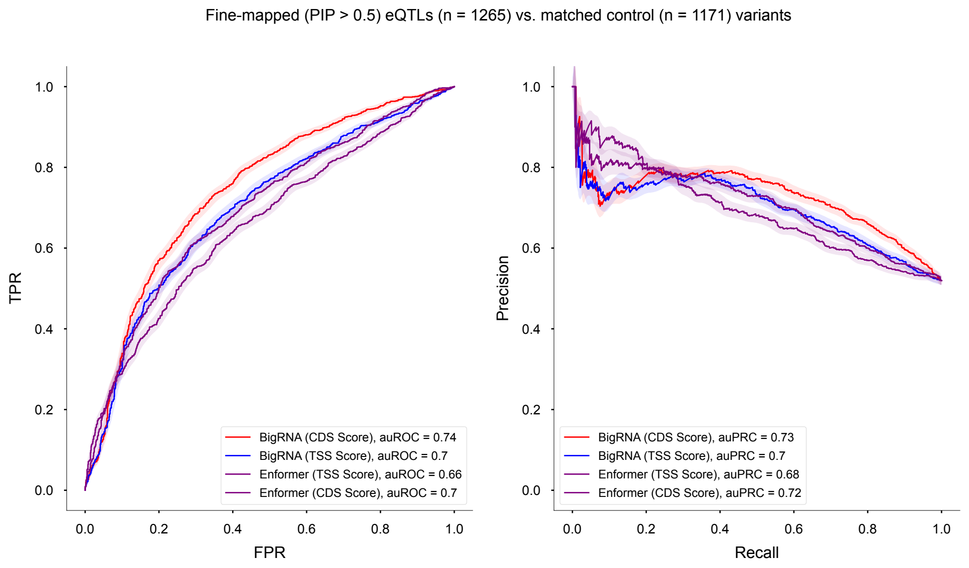

**Supplementary Figure S14** Precision recall and receiver operator characteristic curves for BigRNA and Enformer using either the transcription start site (TSS) interval or the contiguous coding sequence (CDS) disjoint interval for score calculation. The CDS score out-performs the TSS score on AUROC ($p = 0.0017$, $p = 0.0099$ for BigRNA and Enformer respectively) but not on AUPRC ($p = 0.161$, $p = 0.107$ for BigRNA and Enformer respectively) using 10,000 bootstrap resampling iterations to assess significance. These evaluations use the ”max” method for aggregating Ensembles and are tissue specific.

In conclusion, this Supplementary Note described the rationale for constructing scores for our eQTL evaluation. Additionally, the relative performance of the score along three axis was evaluated. From this evaluation it can be concluded that i) the CDS score out-performs the TSS score, ii) scoring in a tissue-specific manner is superior to scoring in a tissue-agnostic manner, and iii) taking the maximum of the Ensemble method scores is a superior approach to taking the mean of the Ensemble method scores. We provide a fully enumerated comparison of all comparisons along three different PIP cut offs (PIP = 0.5, 0.8, and 0.95) and distance thresholds (all, distal (> 10kb), and proximal (< 10kb)) in **Supplementary Data 6**.

[3] Consortium, T.G., Aguet, F., Anand, S., Ardlie, K.G., Gabriel, S., Getz, G.A., Graubert, A., Hadley, K., Handsaker, R.E., Huang, K.H., Kashin, S., Li, X., MacArthur, D.G., Meier, S.R., Nedzel, J.L., Nguyen, D.T., Segrè, A.V., Todres, E., Balliu, B., Barbeira, A.N., Battle, A., Bonazzola, R., Brown, A., Brown, C.D., Castel, S.E., Conrad, D.F., Cotter, D.J., Cox, N., Das, S., Goede, O.M., Dermitzakis, E.T., Einson, J., Engelhardt, B.E., Eskin, E., Eulalio, T.Y., Ferraro, N.M., Flynn, E.D., Fresard, L., Gamazon, E.R., Garrido-Martín, D., Gay, N.R., Gloudemans, M.J., Guigó, R., Hame, A.R., He, Y., Hoffman, P.J., Hormozdiari, F., Hou, L., Im, H.K., Jo, B., Kasela, S., Kellis, M., Kim-Hellmuth, S., Kwong, A., Lappalainen, T., Li, X., Liang, Y., Mangul, S., Mohammadi, P., Montgomery, S.B., Muñoz-Aguirre, M., Nachun, D.C., Nobel, A.B., Oliva, M., Park, Y., Park, Y., Parsana, P., Rao, A.S., Reverter, F., Rouhana, J.M., Sabatti, C., Saha, A., Stephens, M., Stranger, B.E., Strober, B.J., Teran, N.A., Viñuela, A., Wang, G., Wen, X., Wright, F., Wucher, V., Zou, Y., Ferreira, P.G., Li, G., Melé, M., Yeger-Lotem, E., Barcus, M.E., Bradbury, D., Krubit, T., McLean, J.A., Qi, L., Robinson, K., Roche, N.V., Smith, A.M., Sobin, L., Tabor, D.E., Undale, A., Bridge, J., Brigham, L.E., Foster, B.A., Gillard, B.M., Hasz, R., Hunter, M., Johns, C., Johnson, M., Karasik, E., Kopen, G., Leinweber, W.F., McDonald, A., Moser, M.T., Myer, K., Ramsey, K.D., Roe, B., Shad, S., Thomas, J.A., Walters, G., Washington, M., Wheeler, J., Jewell, S.D., Rohrer, D.C., Valley, D.R., Davis, D.A., Mash, D.C., Branton, P.A., Barker, L.K., Gardiner, H.M., Mosavel, M., Siminoff, L.A., Flicek, P., Haeussler, M., Juettemann, T., Kent, W.J., Lee, C.M., Powell, C.C., Rosenbloom, K.R., Ruffier, M., Sheppard, D., Taylor, K., Trevanion, S.J., Zerbino, D.R., Abell, N.S., Akey, J., Chen, L., Demanelis, K., Doherty, J.A., Feinberg, A.P., Hansen, K.D., Hickey, P.F., Jasmine, F., Jiang, L., Kaul, R., Kibriya, M.G., Li, J.B., Li, Q., Lin, S., Linder, S.E., Pierce, B.L., Rizzardi, L.F., Skol, A.D., Smith, K.S., Snyder, M., Stamatoyannopoulos, J., Tang, H., Wang, M., Carithers, L.J., Guan, P., Koester, S.E., Little, A.R., Moore, H.M., Nierras, C.R., Rao, A.K., Vaught, J.B., Volpi, S.: The GTEx Consortium atlas of genetic regulatory effects across human tissues. Science **369**(6509), 1318–1330

(2020) https://doi.org/10.1126/science.aaz1776

[12] Fernandes, R.C., Toubia, J., Townley, S., Hanson, A.R., Dredge, B.K., Pillman,

K.A., Bert, A.G., Winter, J.M., Iggo, R., Das, R., Obinata, D., Sandhu, S., Risbridger, G.P., Taylor, R.A., Lawrence, M.G., Butler, L.M., Zoubeidi, A., Gregory, P.A., Tilley, W.D., Hickey, T.E., Goodall, G.J., Selth, L.A.: Post-transcriptional gene regulation by MicroRNA-194 promotes neuroendocrine transdifferentiation in prostate cancer. Cell Reports **34**(1), 108585 (2021) https://doi.org/10.1016/j.celrep.2020.108585

[21] Farazi, T.A., Hoeve, J.J., Brown, M., Mihailovic, A., Horlings, H.M., Vijver,

M.J., Tuschl, T., Wessels, L.F.: Identification of distinct miRNA target regulation between breast cancer molecular subtypes using AGO2-PAR-CLIP and patient datasets. Genome Biology **15**(1), 9 (2014) https://doi.org/10.1186/gb-2014-15-1-r9

[30] Picard toolkit. Broad Institute (2019)

[31] Ge, X., Chen, Y.E., Song, D., McDermott, M., Woyshner, K., Manousopoulou, A., Wang, N., Li, W., Wang, L.D., Li, J.J.: Clipper: p-value-free FDR control on high-throughput data from two conditions. Genome Biology **22**(1) (2021) https://doi.org/10.1186/s13059-021-02506-9

[32] Manakov, S.A., Shishkin, A.A., Yee, B.A., Shen, K.A., Cox, D.C., Park, S.S., Foster, H.M., Chapman, K.B., Yeo, G.W., Nostrand, E.L.V.: Scalable and deep profiling of mRNA targets for individual microRNAs with chimeric eCLIP (2022) https://doi.org/10.1101/2022.02.13.480296

[33] Liu, L., Jiang, H., He, P., Chen, W., Liu, X., Gao, J., Han, J.: On the variance of the adaptive learning rate and beyond. In: International Conference on Learning Representations (2020)

[34] Kingma, D.P., Ba, J.: Adam: A Method for Stochastic Optimization (2017)

[41] O'Leary, N.A., Wright, M.W., Brister, J.R., Ciufo, S., Haddad, D., McVeigh, R., Rajput, B., Robbertse, B., Smith-White, B., Ako-Adjei, D., Astashyn, A., Badretdin, A., Bao, Y., Blinkova, O., Brover, V., Chetvernin, V., Choi, J., Cox, E., Ermolaeva, O., Farrell, C.M., Goldfarb, T., Gupta, T., Haft, D., Hatcher, E.,

Hlavina, W., Joardar, V.S., Kodali, V.K., Li, W., Maglott, D., Masterson, P., McGarvey, K.M., Murphy, M.R., O'Neill, K., Pujar, S., Rangwala, S.H., Rausch, D., Riddick, L.D., Schoch, C., Shkeda, A., Storz, S.S., Sun, H., Thibaud-Nissen, F., Tolstoy, I., Tully, R.E., Vatsan, A.R., Wallin, C., Webb, D., Wu, W., Landrum, M.J., Kimchi, A., Tatusova, T., DiCuccio, M., Kitts, P., Murphy, T.D., Pruitt, K.D.: Reference sequence (RefSeq) database at NCBI: current status, taxonomic expansion, and functional annotation. Nucleic Acids Research **44**(D1), 733–745 (2015) https://doi.org/10.1093/nar/gkv1189
